# Clustering of activated CD8 T cells around malaria-infected hepatocytes is rapid and is driven by antigen-specific T cells

**DOI:** 10.1101/508796

**Authors:** Reka K. Kelemen, Harshana Rajakaruna, Ian A. Cockburn, Vitaly V. Ganusov

## Abstract

Copyrighted by the AAI (abstract was published as a part of AAI annual meeting).

## 1 Introduction

Malaria is a life-threatening disease that is a result of red blood cell (erythrocyte) destruction by eukaryotic parasites of the *Plasmodium* genus. The majority of deaths (estimated to be about 500,000 annually) are among children, who have not yet developed immunity to the pathogen [1, 2]. There are five species that infect humans: *P. falciparum, P. vivax, P. malariae, P. ovale*, and *P. knowlesi* [3]. Three species of malaria parasites that are used as animal models for human malaria in mice are *P. yeolii, P. berghei*, and *P. chabauidi* [4]. While there are similarities and differences in replication and pathogenesis of Plasmodium species in humans and mice, in this paper we focus solely on infection of mice with Plasmodium parasites.

The infection of the host is started by a mosquito, the vector between mammalian hosts, injecting the sporozoite form of parasites into the skin. Studies have estimated that the initial number of sporozoites entering the host is as low as 10-50 [5, 6], of which only a fraction succeed to migrate to the liver to start an infection of hepatocytes by forming liver stages [7–9]. This liver stage of infection lasts for approximately 6.5 days in humans and about 2 days in mice [10–13]. Because liver stage is asymptomatic, removal of all liver stages prevents clinical symptoms of malaria and thus is highly desirable feature of an effective vaccine. Indeed, previous studies have shown that memory CD8 T cells are required for protection against a challenge with a relatively large number of sporozoites [14, 15] and that vaccination that induces exclusively memory CD8 T cells of a single specificity can mediate sterilizing protection against a sporozoite challenge [16–23]. Antibodies and CD4 T cells may also contribute to protection in some circumstances, for example, following inoculation of sporozoites by mosquitoes in the skin [24, 25]. Given that mouse liver contains about 1 − 2 × 10^8^ hepatocytes [26–28] and only a tiny proportion of these are infected the ability of memory CD8 T cells of a single specificity to locate and eliminate all liver stages within 48 hours is remarkable. Yet, specific mechanisms by which T cells achieve such an efficiency remain poorly defined.

Recent studies utilizing fluorescently labeled sporozoites and activated Plasmodium-specific CD8 T cells and intravital microscopy revealed clustering of CD8 T cells near the parasite in the mouse livers whereby multiple T cells were located in close proximity (≤ 40 *μ*m) of some liver stages [23, 29–31]. Interestingly, we observed that clustering of T cells near the parasite results in a higher chances of parasite’s death suggesting that clusters may increase the efficiency at which T cells eliminate the infection. Recent *in vivo* studies also found that the killing of virus-infected cells occurs faster when several T cells are near an infected cell [32] that is consistent with a previous report estimating that killing of targets in vivo follows the law of mass-action [33] (meaning that the rate of killing is directly proportional to the concentration of the killers and targets).

Clustering of T cells around Plasmodium liver stages in mice was not uniform as the majority of parasites had no T cells around them (at 6 hours after T cell transfer), while some parasites were surrounded by 20-25 T cells [29]. We have developed three alternative mathematical models aimed at explaining this observed variability in cluster formation and by fitting the models to a subset of the data concluded that the data are best explained by a model in which formation of clusters is driven by a positive feedback loop — clusters of a large size attract more T cells to the site of infection [29]. Analysis of T cell movement in the liver suggested that there may be a bias towards the infection site [34]. Additional experiments revealed a significant correlation between the rate at which new T cells locate the infection site and the number of T cells found in the cluster and independence of the per capita rate at which T cells leave the cluster from the cluster size — both observations were consistent with the “density-dependent” recruitment model.

Yet, our previous analysis did not investigate several other important issues of cluster formation. In particular, we did not fully determine the role of the environment in the formation of clusters around Plasmodium liver stages. For example, the observed correlation between entry rate into a cluster and cluster size could simply arise because some parasites may accidentally “attract” more T cells (e.g., due to higher induced inflammation or a higher blood flow rate, [35, 36]). In addition, we have observed that transfer of activated T cells, specific to Plasmodium, and T cells of irrelevant specificity resulted in co-clustering of T cells of two types [29]; however, whether “non-specific” T cells contributed to the formation of clusters was not determined. Finally, our previous analyses did not determine the kinetics of the cluster formation by assuming that cluster size reaches a steady state by the time of imaging. In this paper we re-analyzed some of the previously published data to further define mechanisms by which clusters of activated CD8 T cells around Plasmodium liver stages are formed.

## 2 Materials and methods

### 2.1 Data

In our analyses we used data from previously published work [29, 37]. Data were generated using an experimental system with Py-specific TCR transgenic T cells (PyTCR). For most of our analyses, data were from experiments involving infection of Balb/c mice with variable doses of Plasmodium yoelii (Py) sporozoites, expressing GFP; location of T cells around GFP-expressing liver stages was then visualized using spinning disk confocal microscopy [29]. Following our previous work we consider T cells located within 40 *μ*m distance from the parasite as being close enough to recognize the infection; thus, all T cells within 40 *μ*m from the parasite are called to form a “cluster”. How well the length of 40 *μ*m represents the size of hepatocytes in mice remains unclear. 2D images of mouse hepatocytes suggested the diameter of 40-80 *μ*m [38]; however, measurements of the total volume of mouse hepatocytes of about *V_h_* = 10^4^ *μ*m^3^ [39] suggest a radius 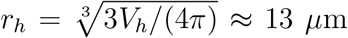 for a “spherical” hepatocyte or cube edge length 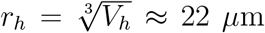 for a “cubical” hepatocyte. A classical textbook on human liver anatomy cites the human hepatocyte volume of 10^4^ – 6 × 10^4^ *μ*m^3^, corresponding to a cube edge of about 40 *μ*m [35, p. 13]. Despite these inconsistent estimates we consider T cells within 40 *μ*m from the parasite to be close enough for recognition of the parasite.

Clustering of CD8 T cells around the parasite was measured in several alternative experiments. In one set of experiments, clustering of CD8 T cells around Py liver stages was performed by immunizing mice with radiation-attenuated Py sporozoites (RAS). In another set of experiments, clustering of T cells was observed following transfer of activated T cell receptor (TCR) transgenic CD8 T cells, specific to epitope located in Py circumsporozoite (CS) protein (CS_280-288_: SYVPSAEQI) denoted as PyTCR cells, or of TCR transgenic CD8 T cells, specific to an epitope in chicken ovalbumin (OVA_257-264_: SIINFEKL) denoted as OT1. Following infection with Py, PyTCR recognize the infection while OT1 cells serve as a control (non-specific to Py) T cells. PyTCR and OT1 cells were activated in similar in vivo experiments (using Vaccinia virus expressing CS or OVA epitopes) [29]. Experiments involving co-transfer of PyTCR and OT1 cells were done in CB6 mice (F1 cross of Balb/c and C57BL/6 mice) [29].

In summary, the following datasets were used in the analysis:

1. Dataset #1. fluorescently labeled PyTCR cells (‘PyTCR alone’) and PyTCR cells pre-treated with pertussis toxin (‘PyTCR+PT’) and transferred into mice infected with Py. This dataset was published before [29] but was not analyzed with mathematical models.
2. Dataset #2: naive or RAS-immunized mice that were infected with Py-GFP and clustering around the parasite was imaged using anti-CD8 antibody. This dataset was published before [29] and was only analyzed with models based on T cell-intrinsic clustering mechanisms (see below).
3. Dataset #3: fluorescently labeled PyTCR and OT1 cells transferred alone into individual mice (‘PyTCR alone’ and ‘OT1 alone’) or as 1:1 mixture (‘PyTCR mix’ and ‘OT1 mix’) to Py-infected mice. This dataset was published before [29] but was analyzed assuming that clustering of PyTCR and OT1 T cells was independent.
4. Dataset #4: fluorescently labeled PyTCR cells transferred into Py-infected mice and imaged at two different time points after T cell transfer. This dataset was generated for a previous study [29] but was not analyzed before with the use of mathematical models.

All datasets are made available as an online supplement to this paper to facilitate further independent analyses.

### 2.2 Mathematical models

#### 2.2.1 Basic mathematical model for clustering of one cell type

Previously we proposed a standard “birth-death” model to describe formation of clusters around Plasmodium liver stages [29]. To be comprehensive, we describe this modeling framework here but then extend it to consider environmental variability in cluster sizes, co-clustering of different T cell specificies, and kinetics of cluster formation. This modeling framework assumes that infection of hepatocytes by Plasmodium sporozoites occurs independently, i.e., there is no interactions between sporozoites infecting different hepatocytes. This assumption of independence is likely to be justified given that in our experiments i) in general ~ 10^5^ sporozoites are injected i.v. into mice, ii) only a fraction of these is expected to reach the liver [7–9, 40, 41], and iii) mouse liver contains 1 – 2 × 10^8^ hepatocytes [26–28]. Because in general in our experiments the number of Plasmodium-specific T cells exceeds the number of liver stages by 10 to 30 fold, we assume that clustering of T cells around one parasite does not interfere or compete with T cells clustering around another parasite. In the model describing formation of clusters around Plasmodium liver stages by T cells of a single specificity we denote *P_k_*(*t*) as the probability to observe *k* T cells around the parasite at time *t* with *k* = 0,1, 2,…*k*_max_. Increase in cluster size occurs at the “birth” (or entry) rate *λ*_*k*_(*k* = 0,1,2,…,*k*_max_) and decline in cluster size occurs due to “death” (or exit) rate *μ_k_*(*k* =1, 2,…, *k*_max_). The mathematical model describing the change in the probability *P_k_*(*t*) with time is given by the system of differential equations:

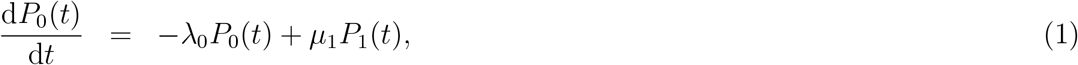

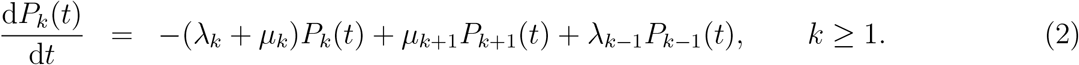

By assuming different specific forms for the T cell entry (λ_*k*_) and exit (*μ_k_*) rates (e.g., see Figure 1 and below) the model can be solved numerically and fitted to the data using maximum likelihood method (see below). For some analyses we made a simplifying assumption that the distribution of cluster sizes reaches a steady state, and the steady state values for the probability to observe *k* CD8 T cells near a given liver stage 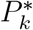 is given by

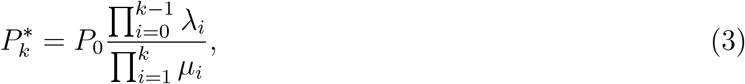

where 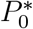 is found by normalizing the total probability to one. By assuming steady state solutions it is in general impossible to estimate individual values for the rates of T cell entry into the cluster and exit from the cluster but we can estimate the ratio of the entry and exit rates, which we define as the relative entry rate *θ_k_* = λ_*k*_/*μ_k_*.

**Figure 1:**
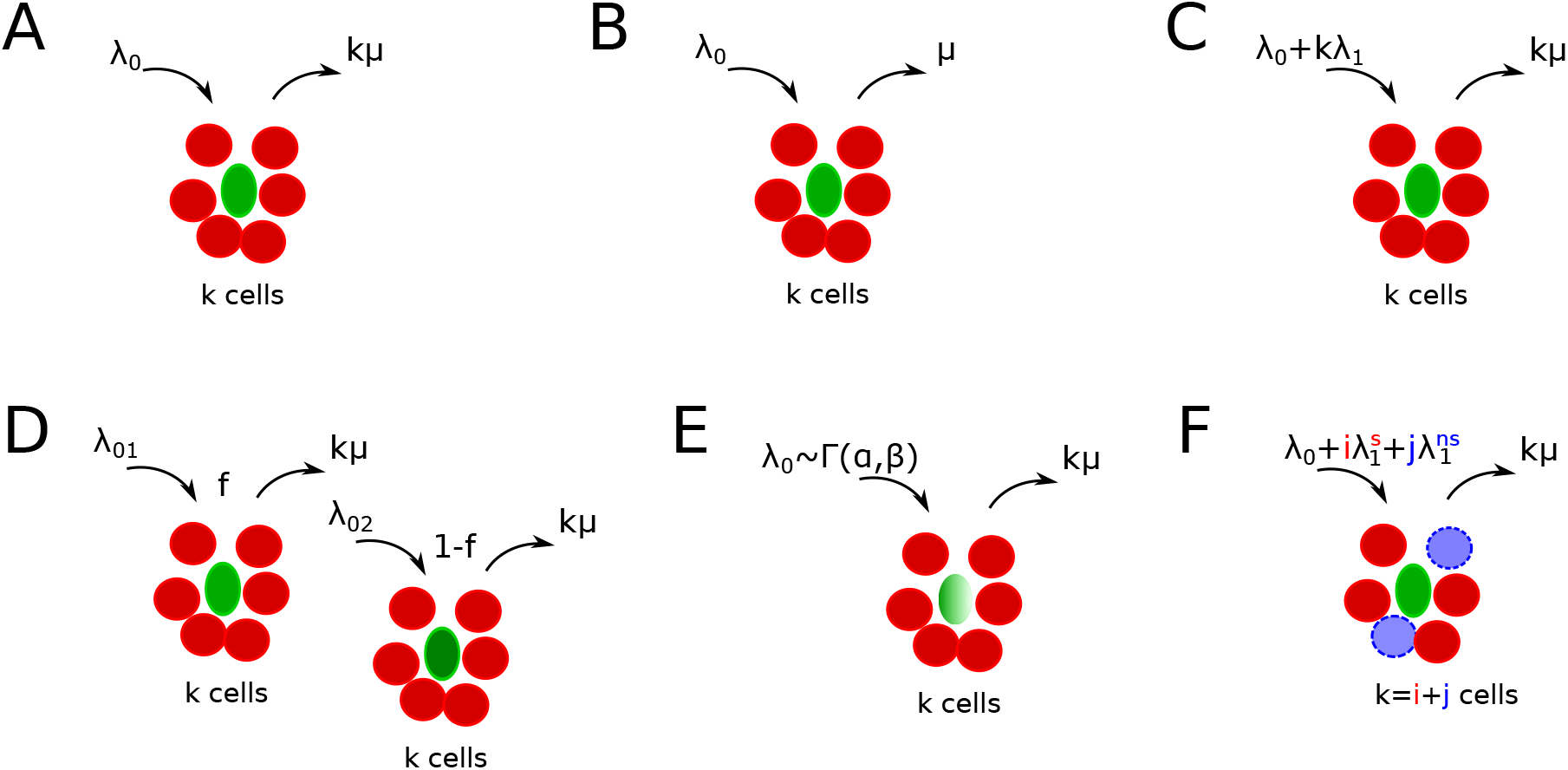
Schematics of alternative mathematical models of T cell cluster formation around Plasmodium yoelii (Py)-infected hepatocytes in mice. Py-specific T cells are labeled by red (discs), T cells of irrelevant specificity are colored by blue (dashed discs), and parasites are labeled by green (ovals). In the models the rate of T cell entry into a cluster is denoted as λ_*k*_ and rate of exit from the cluster is denoted as *μ_k_*. Mathematical models include a random entry/exit (Poisson) model (A, eqn. (4), λ_*k*_ = λ_0_ and *μ_k_* = *kμ*), a density-independent (DI) exit model (B, eqn. (5), λ_*k*_ = λ_0_ and *μ_k_* = *μ*), a density-dependent (DD) recruitment model (C, eqn. (6), λ_*k*_ = λ_0_ + *k*λ_1_ and *μ_k_* = *kμ*), a “two populations” model in which infected hepatocytes have either of two different “attractiveness” levels determined by λ_01_ and λ_02_ (D, eqn. (10), *μ_k_* = *kμ*), a “gamma” model, in which the entry rate into clusters is distributed according to a gamma distribution with *α* and *β* being the rate and shape parameters (E, eqn. (11), *μ_k_* = *kμ*), and finally a “co-clustering” model, in which clusters are formed by Plasmodium-specific T cells or T cells or irrelevant specificity (nonspecific T cells) (F, eqns. (12)–(15), 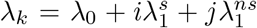 and *μ_k_* = *kμ*). For some of our analyses we characterized the model behavior using the ratio of entry to exit rates denoted as a relative entry rate *θ_k_* = λ_*k*_/*μ_k_*.

Mechanisms explaining the clustering of T cells around Plasmodium parasites in the liver can be broadly divided into two categories: T cell-intrinsic and T cell-extrinsic. In the T cell-intrinsic mechanism, the formation of clusters is driven exclusively by T cells and thus this mechanism ignores any potential differences in the variability in local liver environment. In the T cell-extrinsic mechanism, formation of clusters is driven exclusively due to variability in the liver environment near individual parasites, for example, due to a higher blood flow to some liver stages or a higher degree of inflammation that individual parasites may induce [35, 36]. It is possible that ultimately both mechanisms may contribute to the cluster formation.

#### 2.2.2 Sub-models assuming T cell-intrinsic clustering mechanisms

We consider several alternative models of how T cells may mediate formation of clusters around Plasmodium liver stages in mice (Figure 1A-C). Some of these models have been presented in our previous publication [29] and here are presented again for completeness. Our simplest random entry/exit (**Poisson model**) assumes that entry into the cluster and exit from the cluster occur randomly, i.e., λ_*k*_ = λ_0_ and *μ_k_* = *kμ* where λ_0_ and *μ* are constants (Figure 1A). Solving eqn. (3), the probability to observe *k* T cells around a parasite according to this random entry/exit model is then given by the Poisson distribution:

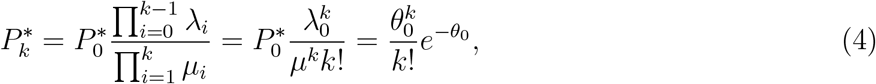

where *θ*_0_ = λ_0_/*μ*.

We have shown previously that the Poisson model is often unable to describe distribution of cluster sizes of Plasmodium-specific CD8 T cells in the liver [29]. One potential mechanism proposed to describe formation of large clusters is a “retention” model in which T cells which recognize the infection, are retained near the parasite. One version of such a model is a **density-independent (DI) exit model** (Figure 1B) in which the rate of T cell exit from a cluster declines with the number of T cells in the cluster, i.e., *μ_k_ = kμ/k = μ* for *k* > 0 and λ_*k*_ = λ_0_ for all *k*. Solving eqn. (3), the probability to observe *k* T cells around a parasite according to the DI exit model is given by a geometric distribution:

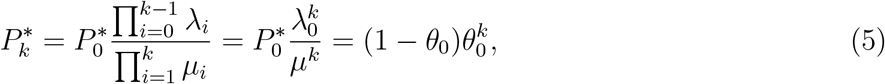

where *θ*_0_ = λ_0_/*μ*. There are other ways in which the total rate of T cell exit from the cluster *μ_k_* could decline with cluster size *k* and in our additional analyses we tested two of such alternative models: a powerlaw model in which *μ_k_* = *k^α^μ* (defined for *k* > 0 with *α* and *μ* being constant) and an exponential model in which *μ_k_* = *kμe*^−*αk*^ (defined for *k* > 0 with α and *μ* being constant). When fitting these retention models to experimental data we did not derive the steady state solutions but instead used numerical solutions of the basic mathematical model (eqns. (1)–(2)).

An alternative mechanism for the formation of large clusters of CD8 T cells around the infection is an “attraction” model in which the rate of T cell entry into the cluster depends on cluster size. In this **density-dependent (DD) recruitment model** (Figure 1C) the entry rate into the cluster is given by λ_*k*_ = λ_0_ + λ_1_*k* while the total exit rate is density-dependent *μ_k_* = *kμ*. Solving eqn. (3), the probability to observe *k* T cells around a parasite according to the DD recruitment model at the steady state is calculated numerically:

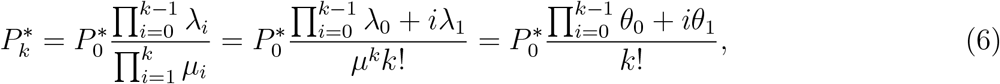

where *θ*_0_ = λ_0_/*μ* and *θ*_1_ = λ_1_/*μ* and 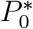 is found by normalizing eqn. (6) assuming the maximal cluster size to be *k*_max_. In general, 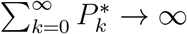 and therefore, the sum must be taken for a finite number of terms due to this reason [29].

To understand dynamics of cluster formation in the Poisson and DD recruitment models it is also useful and possible to derive the model describing the change in the average number of T cells around the parasite (average cluster size), 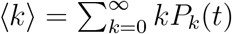 using standard methods of physical chemistry [42]

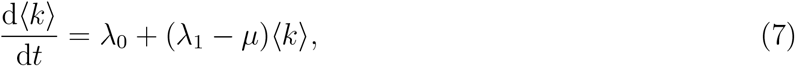

which is a standard birth-death process with immigration which for 〈*k*〉(0) = 0 has the solution

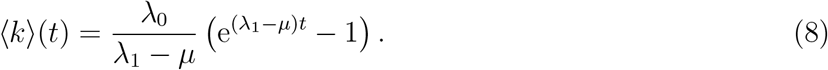

In cases when λ_1_ > *μ* the average cluster size grows indefinitely with time. When λ_1_ < *μ*, which is often found in our analyses (see Main text), average cluster size at the steady state is given by

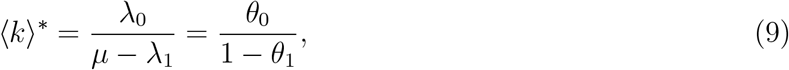

where *θ*_0_ and *θ*_1_ are defined after eqn. (6).

#### 2.2.3 Sub-models assuming T cell-extrinsic clustering mechanisms (environment)

An alternative mechanism for the formation of T cell clusters around Plasmodium-infected hepatocytes is proposed in this paper, namely, that the formation of clusters is driven by the ability of different parasites to “attract” T cells. For example, some parasites while traveling from the blood to hepatocyte or while replicating in the hepatocyte may induce higher degree of inflammation than other parasites, thus, potentially increasing the chance of finding such “inflamed” sites by T cells. Indeed, sporozoites are able to induce inflammation in the liver [36].

We consider two versions of the “environment” model in which T cell recruitment to sites is determined by the variability in parasite’s “attractiveness”. In one such version, a **two population model**, we assume that there are parasites of two types found at frequencies *f* and 1 – *f*, and these parasites differ in the rate at which T cells find them (Figure 1D). The formation of clusters around parasites of a given parasite type is given by random entry/exit model with rates λ_01_ and λ_02_ while the rate of exit of T cells from the cluster is *μ_k_* = *kμ*. Then assuming a steady state the probability to observe clusters of size *k* is given by

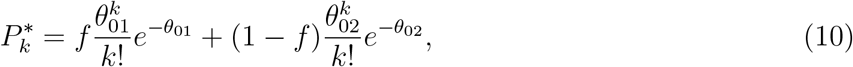

where *θ*_01_ = λ_01_/*μ* and *θ*_02_ = λ_02_/*μ*. Alternatively, the rate at which T cells find parasites could be given by a continuous function, and we tested a model in which entry rate into the cluster is given by a gamma distribution 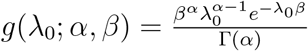, i.e., the probability for T cells to have an entry rate in the interval (λ_0_, λ_0_ + dλ_0_) is *g*(λ_0_; *α, β*)dλ_0_. The probability to observe a cluster of size *k* given that clustering around a parasite “attracting” T cells at a rate λ follows a Poisson model is given by an integral

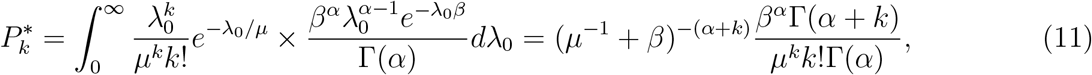

where *α* and *β* are the shape and rate parameters of the Gamma distribution, respectively, and Γ(*α*) = (*α* − 1)!.

#### 2.2.4 A basic mathematical model for clustering of two cell types

In some of our experiments we tracked clustering of T cells of two specificities: one type of T cells was specific to Plasmodium sporozoites (PyTCR) and another type of T cells was specific to irrelevant antigen (OT1). To quantify the kinetics of clustering of Plasmodium-specific (PyTCR) and nonspecific T cells (OT1) around Plasmodium-infected hepatocytes, we extended our basic model (eqns. (1)–(2)) to include two types of cells, *t*_1_ and *t*_2_, in the cluster to formulate a **co-clustering model** (Figure 1F). We define *P_ij_*(*t*) as the probability to observe *i* cells of type 1 and *j* cells of type 2 in a given cluster. Then the rate at which new T cells of type *x*, where *x* = *t*_1_, *t*_2_ enter the cluster with *i* T cells of type 1 and *j* T cells of type 2 is 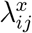. Similarly, 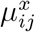 is the rate of exit of T cell of type *x* from a cluster with (*i, j*) T cells. The dynamics of the probability to observe a cluster with (*i, j*) T cells is given by equations

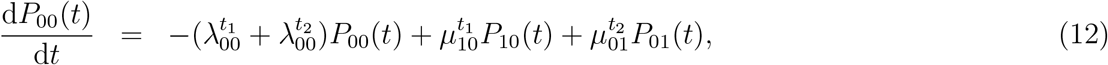

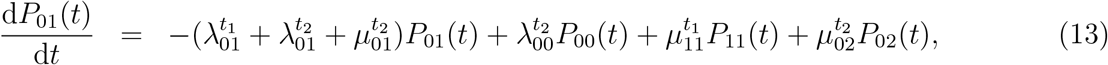

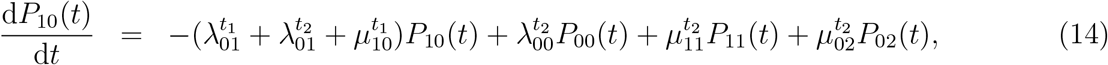

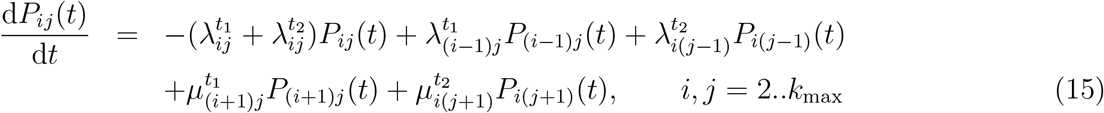

The dynamics of the probability *P_ij_*(*t*) can be simulated by assuming different functional forms for the entry and exit rates (Table 1). For example, when the entry rate into the cluster is independent of the cluster size or cell type, 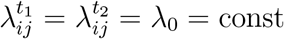, and the exit rate is dependent on the number of T cells of a given specificity present near the parasite, 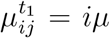 and 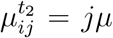 where *μ* = const, clustering of T cells is independent and is described at the steady state by the Poisson distribution (results not shown). Another model is when the entry rate of T cells into the cluster is dependent only on the number of specific T cells (*t*_1_) already in the cluster: 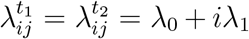 with exit rates being similar to the random entry/exit model described above.

**Table 1:**
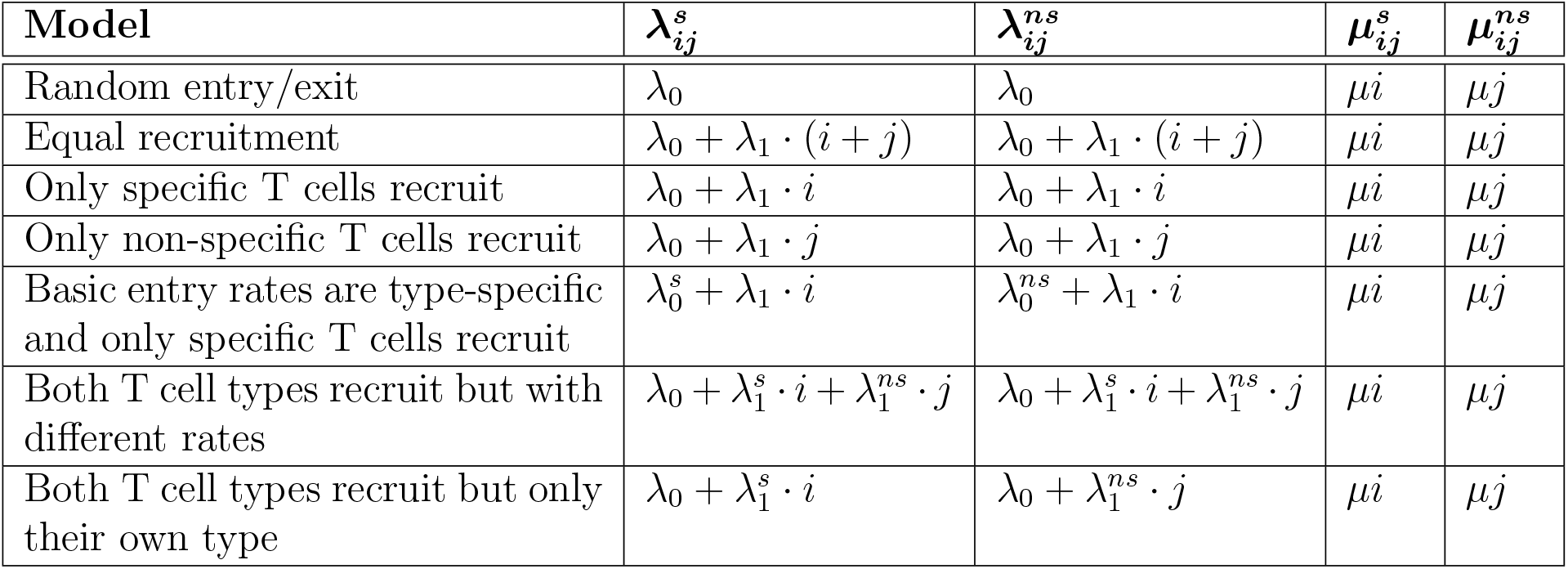
Defining alternative models for co-clustering of Plasmodium-specific (*s*) and non-specific (*ns*) T cells around Plasmodium liver stages. For the general mathematical model describing co-clustering of two cell types (eqns. (12)–(15)) we define parameters determining the rate of T cell entry into the cluster (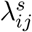 and 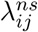) and the rate of exit from the cluster (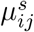 and 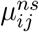) where superscripts “*s*” and “*ns*” stand for Plasmodium-specific and non-specific T cells, respectively, *ij* denotes a cluster with *i* specific and *j* nonspecific T cells around a given liver stage. For example, 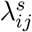 denotes the entry rate of specific T cells into a cluster with *i* specific and *j* non-specific T cells. Parameters λ_0_ (the initial entry rate), *μ* (per capita exit rate), and λ_1_ (increase in entry rate with cluster size) are found by fitting the numerical solution of the mathematical model (given in eqns. (12)–(15)) to the co-clustering data (dataset #3).

| Model | $\lambda_{ij}^s$ | $\lambda_{ij}^{ns}$ | $\mu_{ij}^s$ | $\mu_{ij}^{ns}$ |
| --- | --- | --- | --- | --- |
| Random entry/exit | $\lambda_0$ | $\lambda_0$ | $\mu i$ | $\mu j$ |
| Equal recruitment | $\lambda_0 + \lambda_1 \cdot (i + j)$ | $\lambda_0 + \lambda_1 \cdot (i + j)$ | $\mu i$ | $\mu j$ |
| Only specific T cells recruit | $\lambda_0 + \lambda_1 \cdot i$ | $\lambda_0 + \lambda_1 \cdot i$ | $\mu i$ | $\mu j$ |
| Only non-specific T cells recruit | $\lambda_0 + \lambda_1 \cdot j$ | $\lambda_0 + \lambda_1 \cdot j$ | $\mu i$ | $\mu j$ |
| Basic entry rates are type-specific and only specific T cells recruit | $\lambda_0^s + \lambda_1 \cdot i$ | $\lambda_0^{ns} + \lambda_1 \cdot i$ | $\mu i$ | $\mu j$ |
| Both T cell types recruit but with different rates | $\lambda_0 + \lambda_1^s \cdot i + \lambda_1^{ns} \cdot j$ | $\lambda_0 + \lambda_1^s \cdot i + \lambda_1^{ns} \cdot j$ | $\mu i$ | $\mu j$ |
| Both T cell types recruit but only their own type | $\lambda_0 + \lambda_1^s \cdot i$ | $\lambda_0 + \lambda_1^{ns} \cdot j$ | $\mu i$ | $\mu j$ |

#### 2.2.5 Stochastic simulations

We simulated cluster formation using the Gillespie algorithm as previously described [43]. In short, for every iteration we first determined randomly the time of the change in cluster size which is determined by the total rate at which clusters could increase or decrease in size (e.g., in the DD recruitment model this rate for a cluster of size *k* is λ_0_ + *k*λ_1_ + *kμ*). The second step was to then choose at random which of two events (cluster size increase or decrease) occurs; this is determined by the relative value of the entry rate into the cluster (e.g., λ_0_ + *k*λ_1_) or exit from the cluster (e.g., *kμ*).

#### 2.2.6 Statistics

Our clustering data are given as the number of T cells found in the 40 *μ*m radius from a given parasite following intravital microscopy imaging [29], i.e., the data are simply a column of integers representing T cell numbers per parasite. (In co-clustering experiments the data also represent the number of Plasmodium-specific and non-specific T cells found per parasite.) As the data shows, in many cases majority of parasites have no T cells associated with them [29, and see main text].

To estimate parameters of mathematical models we used a likelihood approach where the likelihood represents the product of probabilities to observe clusters of different sizes

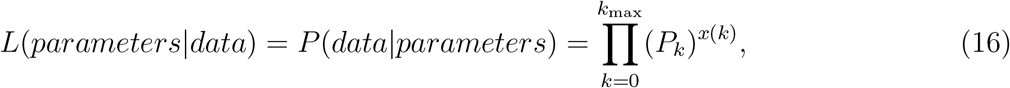

where *P_k_* is the mathematical model-predicted probability of observing a cluster of size *k* according to a set of parameter values, *x*(*k*) is the number of clusters of size *k* in the data, and *k*_max_ is the maximal cluster size in the data. In this procedure, the probability *P_k_*(*t*) can be either given analytically as a steady state solution (e.g., eqn. (4)) or can be found by numerically solving the basic mathematical model predicting *P_k_*(*t*) at a particular time (e.g., eqns. (1)–(2)). When fitting numerical solutions of the model to experimental data in some cases we fixed the rate of exit of T cells from clusters μ to different values because we found that it is generally impossible to accurately estimate both entry and exit rates simultaneously (see Results section).

The models were fitted by calculating negative log-likelihood 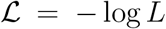 and using routine `FindMinimum` in `Mathematica` version 11. When alternative models were fitted to the same dataset, we compared quality of the model fits to the data by comparing Akaike weights *w* based on the corrected Akaike Information Criterion (AIC) [44]:

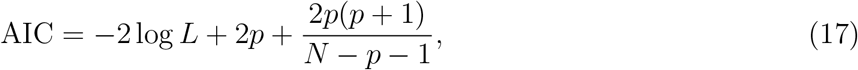

where *p* is the number of model parameters and *N* is the number of data points (parasites). AIC provides a score for each model based on its maximum likelihood value and the number of model parameters. The model with the lowest AIC score is considered to be best relative to the tested models. Weights of a given model can be treated as a likelihood of the model in the list of the tested models. As a rule of thumb, models with *w* < 0.05 can be considered to be inconsistent with experimental data in favor for models with higher weights. In addition, when comparing nested models, where one model is a special case of another, we used the likelihood ratio test.

## 3 Results

### 3.1 Pertussis toxin-treated PyTCR cells form clusters randomly

By using intravital microscopy two recent studies have observed that activated CD8 T cells, specific to Plasmodium antigens, often localize near Plasmodium liver stages [29, 30]. In our experiments we used GFP-expressing *Plasmodium yoelii* (Py) sporozoites and fluorescently-labeled activated Py-specific CD8 T cells (PyTCR cells) [29]. A peculiar feature of these observations was a variable number of T cells found near individual parasites: in many experiments multiple parasites had no T cells while some parasites had more than 20 T cells in a 40 *μ*m radius [29]. We have developed three alternative mathematical models that assume that the formation of clusters of CD8 T cells around Plasmodium liver stages is T-cell-intrisitic and is mediated exclusively by T cells. These models include 1) the null model predicting formation of clusters by chance (“random entry/exit” or Poisson model, eqn. (4)), 2) a model in which T cells find parasites randomly but are retained near the parasite (“density-independent (DI) exit” model, eqn. (5)), and 3) a model in which T cells actively recruit other T cells to the infection site (“density-dependent (DD) recruitment” model, eqn. (6)). By fitting these models to experimental data on clustering we concluded that the DD recruitment model is best consistent with data on clustering of PyTCR cells [29]. By performing experiments in which either PyTCR cells or PyTCR cells, treated with pertussis toxin (PyTCR+PT cells), were transferred into mice, previously infected with Py, we found that PT-treated PyTCR cells form smaller clusters that untreated T cells (Figure 2). PT treatment inactivates G-protein-coupled receptors (GPCRs) in the cells, and in particular, makes PT-treated cells to be unresponsive to chemokine gradients [45]. Fitting three basic mathematical models, predicting steady state distribution of T cells around the liver stages, to the data on clustering of PyTCR cells (see Materials and Methods for more detail on how models were fitted to data) confirmed our previous result that DD recruitment model provided the best fit of clustering data (Figure 2B). The random entry/exit model did not fit these data adequately, while the DI exit model provided a visually reasonable fit which was not supported by the Akaike weights or log-likelihood (result not shown). Interestingly, plotting the data and model fits on the log-scale revealed that even the best fit model did not accurately predict formation of a large cluster with 21 PyTCR cells (Figure 2B). Indeed, comparing the prediction of the DD recruitment model with data using a goodness-of-fit *χ*^2^ test showed that the model described the clusters until size 8 well (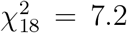, *p* = 0.99), but failed at describing all clusters including one with 21 cells (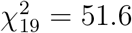, *p* < 0.001). According to the prediction of the mathematical model, the average cluster size of T cells around the liver stages at the steady state is rather small, 〈*k*〉* ≈ 1.5 (see eqn. (9)), because most parasites were not found by T cells within 6 hours (Figure 2A).

**Figure 2:**
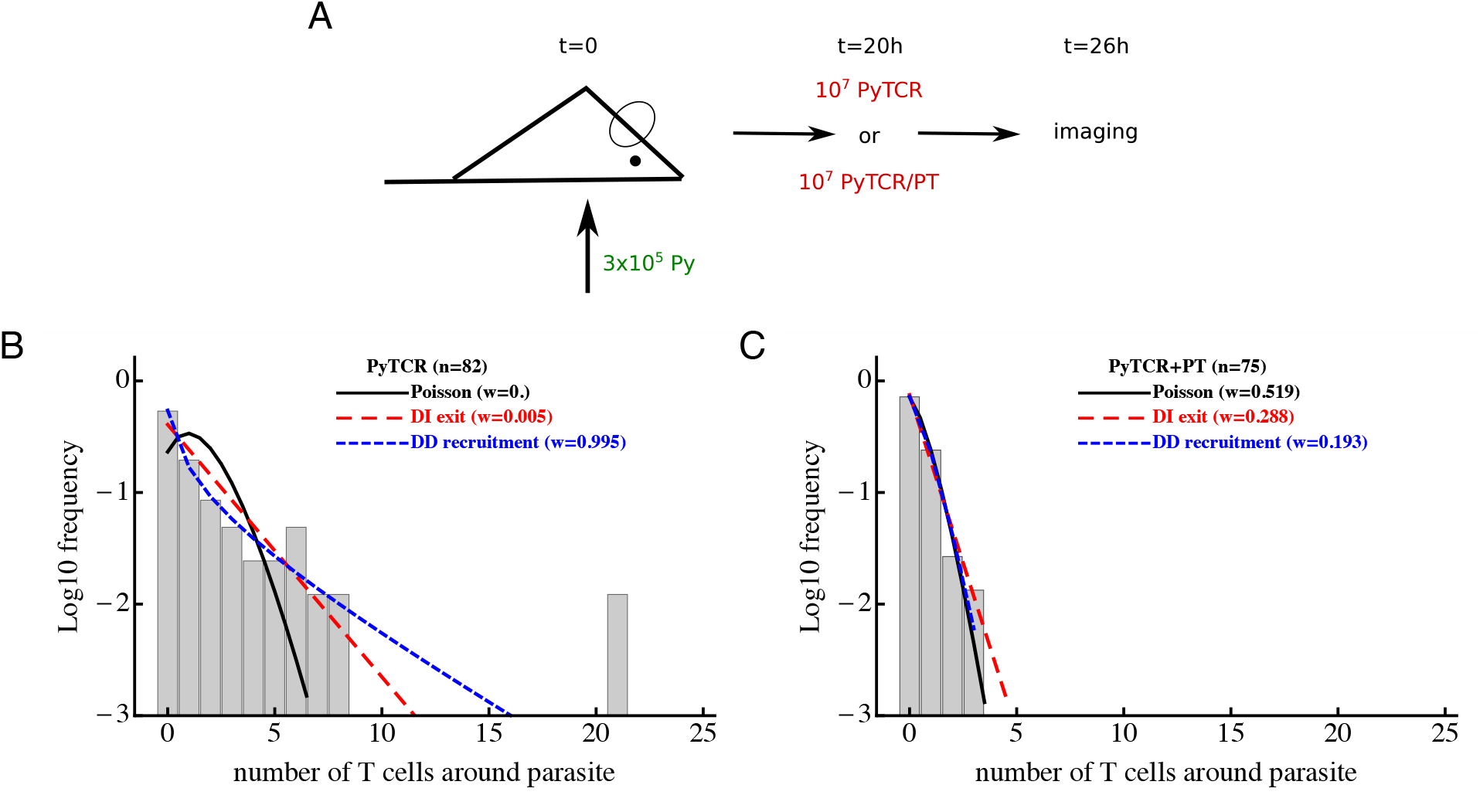
Pertussis toxin (PT)-treated T cells form clusters around Plasmodium yoelii (Py)-infected hepatocytes randomly. Panel A: mice were infected with 3 × 10^5^ GFP-expressing Py sporozoites. Twenty hours later 10^7^ Py-specific activated CD8 T cells (PyTCR) or 10^7^ PyTCR T cells pretreated for one hour with 1 *μ*g/mL PT were transferred into infected mice and imaged with intravital microscopy 6 hours later [29]. The number of T cells in 40 *μ*m radius of randomly chosen n = 82 (PyTCR) or *n* = 75 (PyTCR+PT) parasites was recorded; the frequency of size of such clusters is shown in panels B-C by bars. Panels B and C: we fitted three mathematical models (Poisson (eqn. (4)), DI exit, (eqn. (5)), and DD recruitment (eqn. (6))) to these experimental data using likelihood approach (eqn. (16)) and calculated Akaike weights (*w*) (see Materials and Methods for more detail). Results suggest that clustering of PyTCR T cells is best described by density-dependent recruitment model while clustering of PT-treated T cells occurs mainly randomly. Parameter estimates of the best fit model and 95% CIs obtained by resampling the data with replacement 10^3^ times in panel B are *θ*_0_ = 0.31 (0.20 – 0.48) and *θ*_1_ = 0.79 (0.59 – 0.88) (DD recruitment model) and in panel C are *θ*_0_ = 0.33 (0.21 – 0.48) (Poisson model).

Importantly, however, all three models could accurately describe the formation of clusters by PT-treated PyTCR cells (Figure 2C) suggesting that the formation of these clusters is most likely due to a random encounter between T cells and the Py-infected hepatocyte. Indeed, the estimated relative entry rate *θ*_0_ was similar for two datasets (see legend of Figure 2) and was only slightly higher than the relative rate estimated in our previous work from other experiments involving co-transfer of PyTCR and OT1 cells [29, see Materials and Methods and below]. Thus, this result further supports the conclusion that GPCR-mediated signaling is important in the formation of CD8 T cell clusters around Plasmodium liver stages.

### 3.2 Clustering of endogenous CD8 T cells does not allow to discriminate between T-cell-intrinsic and T-cell-extrinsic models of cluster formation

Our previous analyses so far attempted to explain mechanisms behind the formation of clusters around Plasmodium liver stages from the T-cell-centric point of view; namely, we assumed that cluster formation is dependent on the presence of T cells (e.g., DD recruitment model, see eqn. (6)). However, it is possible that a very different alternative mechanism drives the formation of clusters, which is T-cell-extrinsic. In this case, variable clustering of T cells around the liver stages is driven by variability in the environment, for example, due to the level of “attractiveness” of individual parasites. This may arise because individual parasites may induce different degrees of inflammation as they migrate from the blood into the liver parenchyma, or some parasites may infect hepatocytes which are located in liver parts with a larger blood flow, increasing the chance of T cells to locate such parasites [35, 46].

To investigate whether a T-cell-extrinsic mechanism can be sufficient to explain the formation of clusters in our experiments we formulated two alternative mathematical models predicting the formation of clusters of different sizes: in the first model we assumed that there are two populations of parasites with different levels of attractiveness/rate of entry λ_01_ and λ_02_ (“2 population” model, Figure 1D and eqn. (10)), and in the second model we assumed that there is a distribution in the level of attractiveness of parasites given by a continuous Gamma distribution (“gamma” model, Figure 1E and eqn. (11)). To test whether models assuming T-cell-intrinsic or T-cell-extrinsic mechanisms of cluster formation perform better, we fitted the models to previously published data on clustering of endogenous CD8 T cells around Py liver stages in mice [29]. In these experiments, mice were left naive or were immunized with radiation-attenuated sporozoites (RAS) and then 10 days later infected with wild-type Py expressing GFP (Figure 3A). Clustering of CD8 T cells around GFP-expressing liver stages was visualized by injecting CD8-binding antibody. We fitted five mathematical models to these data using a likelihood approach (eqn. (16)). This analysis showed that all mathematical models could accurately describe the lack of formation of large (*k* > 5) clusters around Plasmodium liver stages in naive (unimmunized) mice, and the simplest (null) random entry/exit model was favored by the Akaike weights (Figure 3B). While all models provided similar likelihood of the model given the data, lower weights for 2 population and gamma models was due to a larger number of fitted parameters (3 in 2 population/gamma models vs. 1 in the Poisson model).

**Figure 3:**
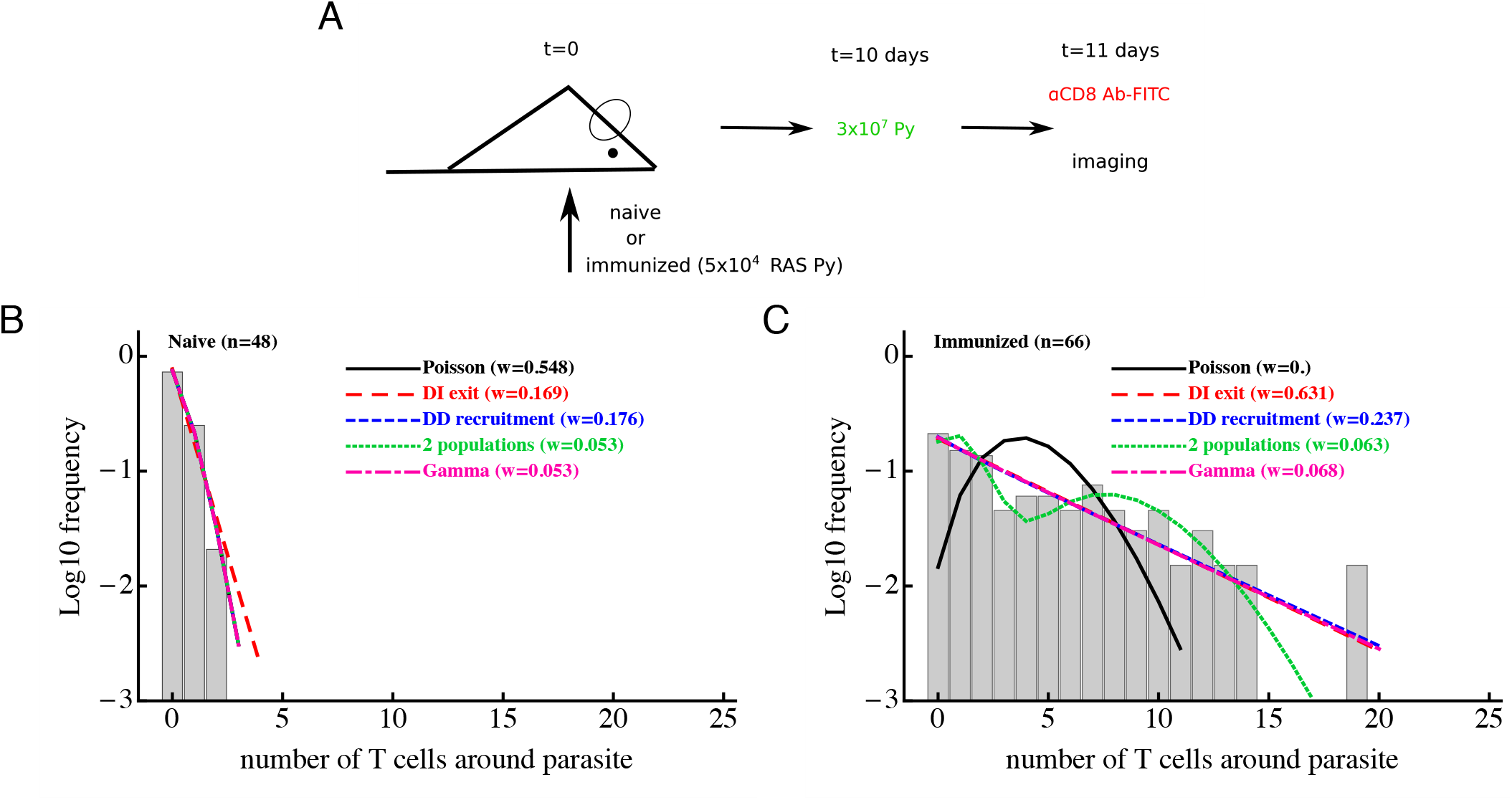
Models assuming time-invariant but spatially-variable environment are consistent with the data on clustering of CD8 T cells in mice immunized with radiation-attenuated sporozoites (RAS). Panel A: mice were immunized with 5 × 10^4^ Py RAS or left unimmunized. Ten days later, mice were infected with 3 × 10^7^ wild-type Py, expressing GFP. One day later CD8 T cells were labeled with 4 *μ*g PE-conjugated anti-CD8 mAbs and clustering of CD8 T cells around Py-infected hepatocytes in the liver was imaged using intravital microscopy [29]. In total 48 (in naive mice, panel B) and 66 (in RAS-immunized mice, panel C) parasites were randomly chosen and the number of T cells in a 40 *μ*m radius were counted. Five different mathematical models were fitted independently to the data on T cell clustering in naive and immunized mice, and the quality of the model fits was evaluated using Akaike weights (*w*). Clustering in naive mice is most consistent with the Poisson (random entry/exit) model, while in RAS-immunized mice models assuming constant environment (“2 populations” and “gamma” models) fit the data worse than other models, in part due to a larger number of parameters than in the DD recruitment or DI exit models. Parameter estimates of the best fit model and 95% CIs in panel B are *θ*_0_ = 0.29 (0.17 – 0.42) (Poisson model) and in panel C are *θ*_0_ = 0.81 (0.77 – 0.84) (DI exit model) or *θ*_0_ = 0.80 (0.52 – 1.16) and *θ*_1_ = 0.82 (0.73 – 0.88) (DD recruitment model). According to the DD recruitment model, the average cluster size in RAS-immunized mice at steady state (panel B) is 〈*k*〉* ≈ 4.2.

Consistent with our previous analysis [29], the random entry/exit (Poisson) model could not adequately describe the distribution of cluster sizes in RAS-immunized mice (Figure 3C). Interestingly, both 2 population and gamma models did not provide an improved fit of these clustering data as compared to DD recruitment or DI exit models which fitted the data with similar quality (Figure 3C). This was surprising given that the DD recruitment and DI exit models were not able to accurately describe the two peaks in the cluster size distribution (at 0 and 7 T cells/parasite). A closer inspection revealed that the DD recruitment, DI exit, 2 population, and gamma models provided fits of nearly identical quality as based on the negative log-likelihood values 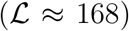, and lower weights were selected for models with more parameters. All models except the Poisson and 2 population models could accurately describe the data (based on goodness-of-fit *χ*^2^ test); the 2 population model deviation was due to its inability to accurately predict the formation of one cluster with 19 cells (results not shown). Importantly, the 2 population and gamma models could fit other clustering data relatively well, e.g., data on clustering of PyTCR cells or PyTCR cells treated with PT (Figure 2, results not shown). Taken together, these results demonstrate that some clustering datasets do not allow to discriminate between T-cell-intrinsic and T-cell-extrinsic mechanisms of formation of CD8 T cell clusters around Plasmodium liver stages.

### 3.3 Environmental variability is not the main driver of cluster formation

To discriminate between T-cell-intrinsic and T-cell-extrinsic mechanisms of formation of CD8 T cell clusters around Plasmodium liver stages we turned to additional experimental data generated previously [29]. In these experiments, Py-specific T cells and T cells of irrelevant specificity (OT1) were transferred either separately or together into mice previously infected with Py-GFP, and the formation of clusters around Py liver stages was measured by intravital microscopy (Figure 4A). We have previously shown that the DD recruitment model describes best (based on Akaike weights) the data on the clustering of PyTCR cells when transferred alone or data on the clustering of PyTCR and OT1 cells when transferred together [29]. In contrast, the clustering of OT1 cells alone was best described by the random entry/exit model [29, Figure 4B-C]. Therefore, these data indicate that the clustering of T cells, which are not specific to Plasmodium depends on the presence of Py-specific T cells suggesting that variability in parasite’s “attractiveness” alone cannot explain these data. We formally tested if the 2 population or gamma models can describe the clustering data of OT1 cells in the following way. We fitted the 2 population model to the data on clustering of OT1 cells alone or in the mixture with PyTCR cells simultaneously. We therefore fitted the models by allowing all three parameters of the model (*θ*_01_, *θ*_01_, and *f*, see eqn. (10)) to be different for the two datasets or by allowing only the fraction of parasites with different attractiveness level *f* to vary between two datasets while keeping other parameters the same between datasets (Figure 4B-C). In this way, we tested the hypothesis that the clustering of OT1 cells is driven exclusively by factors which are independent of Py-specific CD8 T cells. Because two fits are from nested models, comparing the quality of the fits revealed that the model assuming PyTCR-cell-independent environment fits the two datasets significantly worse (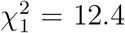, *p* < 0.001). Fitting the gamma model to the same two datasets assuming either identical or variable parameters between the two datasets also suggested that the model with constant parameters fits the data significantly worse (results not shown). Thus, these results strongly suggest that the T-cell-extrinsic models of cluster formation are not consistent with the data on different clustering patterns of OT1 cells in the absence or presence of Py-specific T cells.

**Figure 4:**
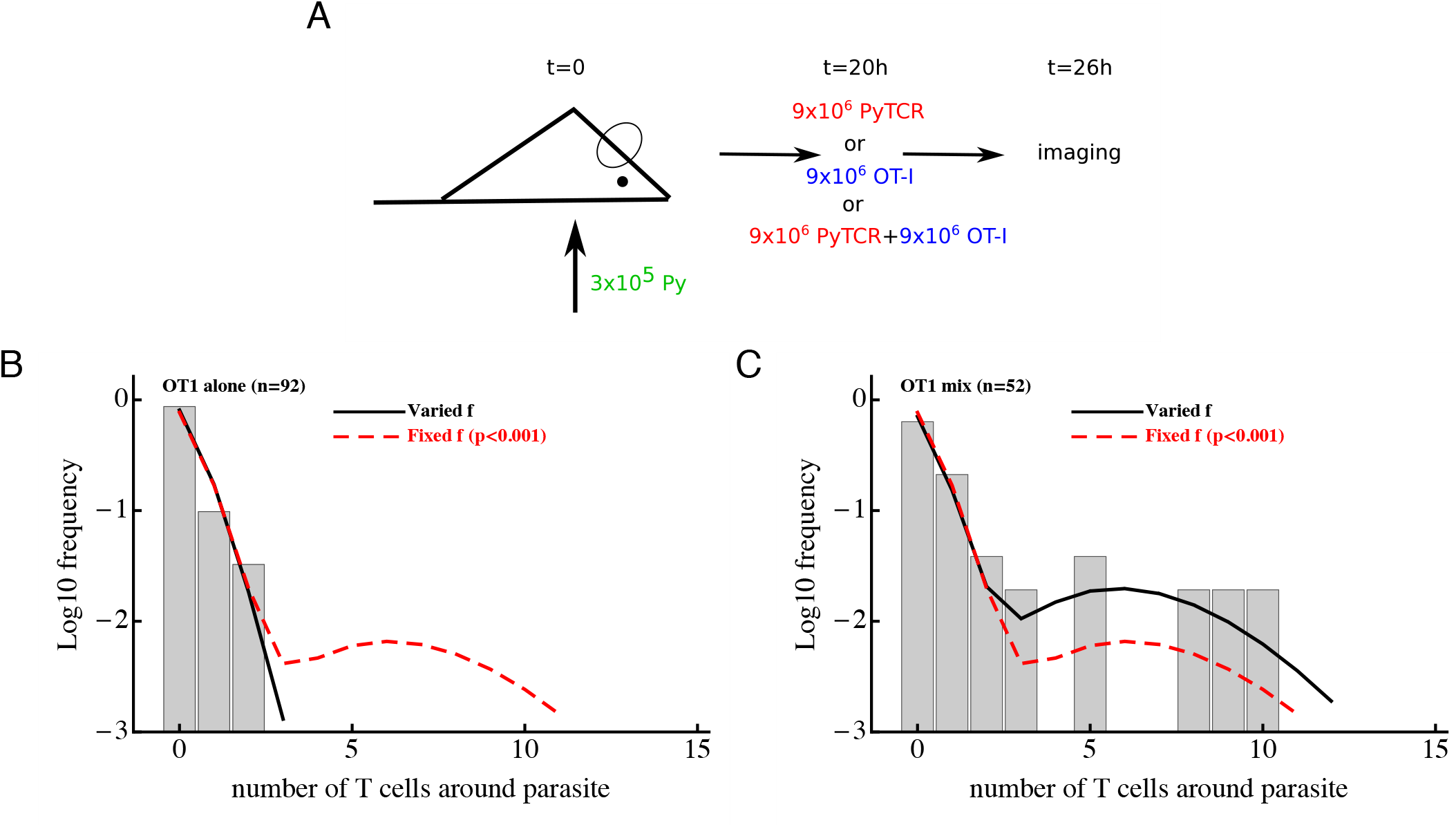
Models assuming time-invariant but spatially-variable environment are unable to accurately describe the clustering of T cells of irrelevant specificity in different conditions. Panel A: mice were infected with 3 × 10^5^ GFP-expressing Py sporozoites. Twenty hours later 9 × 10^6^ Py-specific activated CD8 T cells (PyTCR), 9 × 10^6^ OT1 T cells (specific to chicken ovalbumin), or mixture of 9 × 10^6^ PyTCR and 9 × 10^6^ OT1 T cells were transferred into infected mice and livers of these mice were imaged using intravital microscopy 6 hours later. In total 92 (mice receiving only OT1 cells, panel B) and 52 (in mice receiving a mix of PyTCR and OT1 cells, panel C) parasites were randomly chosen and the number of T cells in a 40 *μ*m radius were counted [29]. The “two population” mathematical model (eqn. (10)) was fitted to these two datasets simultaneously assuming two different entry rates *θ*_01_ and *θ*_02_ and either allowing the fraction of attracting parasites *f* to vary between the datasets (solid line) or to be fixed between the datasets (dashed line). Fixing the fraction *f* between the datasets significantly reduced the quality of the model fit of the data as compared to the model in which *f* could vary (likelihood ratio test, 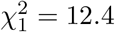, *p* < 0.001).

It is important to note that the use a specific mathematical model (e.g., 2 population model) simply allows to formally test if distributions of cluster sizes of OT1 cells are different in two different conditions. This can be also done using a *χ*^2^ test [47] which showed that these distributions are only marginally different (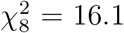, *p* = 0.04). Thus, the use of models allows to obtain much stronger statistical power at falsifying the T-cell-independent (“environment”) hypothesis as the sufficient mechanism of cluster formation.

### 3.4 Several alternative retention models poorly describe data on clustering of PyTCR cells

While our experiments of clustering of OT1 T cells either alone or in presence of PyTCR T cells argue against T-cell-extrinsic clustering model, they do not allow to fully discriminate between alternative T-cell-intrinsic clustering models. Fitting the steady state prediction of the DI exit and DD recruitment model to clustering of PyTCR T cells (Figure 2) or clustering of OT1 T cells in the presence of PyTCR cells [29] favored the DD recruitment model (based on Akaike weights). However, it is possible that a specific form of the retention model, i.e., that the per capita exit rate is inversely proportional to the cluster size, was an accidentally poor choice. Therefore, we tested two alternative models of how exit rate from a cluster could depend on cluster size with *μ_k_* = *k^α^μ* or *μ_k_* = *kμ*e^−αk^. We fitted the numerical solution of the basic mathematical model (eqns. (1)–(2)) to the clustering of PyTCR T cells (Figure 2A) using a likelihood approach. Both alternative retention models still described the data worse than the DD recruitment model (*w* < 0.001 and *w* = 0.02 for the two models, respectively, results not shown) suggesting limited support for the hypothesis that retention of T cells plays the major role in cluster formation. Therefore, in our following analyses we focus exclusively on the DD recruitment model.

### 3.5 No evidence that T cells of irrelevant specificity influence clustering

In our previous analysis we showed that the DD recruitment model-based fit of the data on the clustering of PyTCR and OT1 cells in the co-transfer experiments (Figure 4A) predicted similar relative recruitment rate parameters *θ*_0_ and *θ*_1_ (see Table S1 in Cockburn *et al*. [29]). However, the previous analysis treated clustering of PyTCR and OT1 cells in the co-transfer experiments independently, and here we extend this analysis by considering potential mechanisms behind coclustering of these two cell populations. First, we found that there was no significant difference in the number of PyTCR or OT1 T cells clusters around Plasmodium liver stages with similar proportions of parasite having more PyTCR or OT1 cells (Figure 5A). To investigate whether the data on the co-clustering of T cells may provide evidence of OT1 T cells assisting in cluster formation we developed a mathematical model tracking the dynamics of co-clustering of two types of cells (see eqns. (12)–(15) in Material and Methods) and fitted that model to the co-clustering data (dataset #3) using a likelihood approach. As we show in the next section, our clustering data do not allow to identify both the rate of T cell entry into the cluster and exit rate from the cluster from measuring clusters at one time point. Therefore, in this analysis we fixed the per capita exit rate *μ* = 0.5/h and estimated entry rates. Our overall results were robust to several other tested values of the exit rate such as *μ* = 0.1/h or *μ* = 3/h even though estimates of the entry rates were strongly dependent on the assumed exit rate (results not shown and see next section).

**Figure 5:**
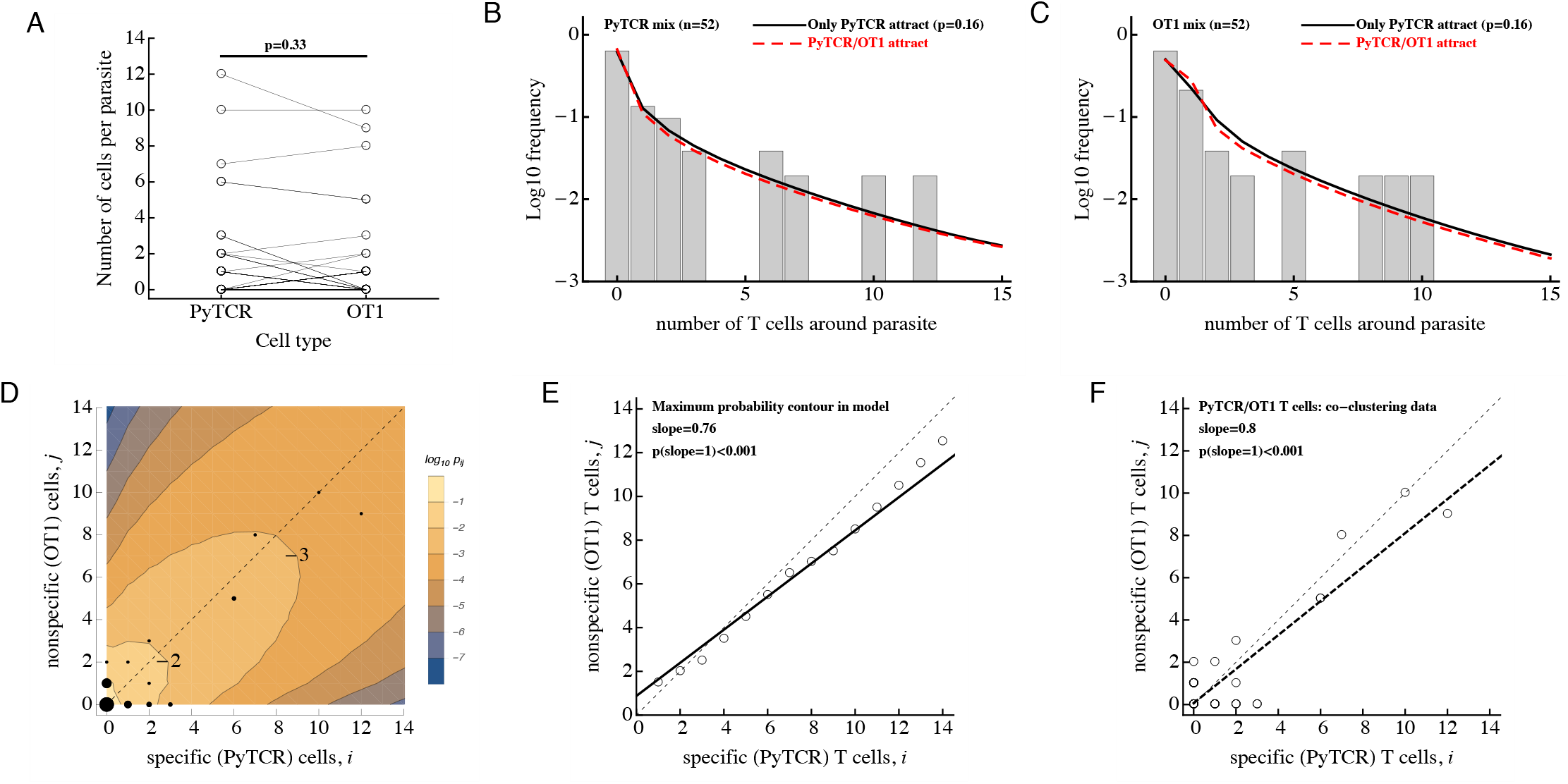
No evidence that activated CD8 T cells of irrelevant specificity play a significant role in cluster formation. Experiments were performed as described in Figure 4A and the number of T cells, specific to Py (PyTCR) and T cells of irrelevant specificity (OT1), found in a 40 *μ*m radius of *n* = 52 randomly chosen parasites in the liver was counted using intravital imaging. Panel A: no difference in the number of Py-specific and non-specific T cells found around Py liver stages (Wilcoxon signed rank test). Panels B-C: we fitted a series of mathematical models assuming how Py-specific or nonspecific T cells mediate attraction to the infection site (co-clustering model, eqns. (12)–(15)), and fits of two models where either only PyTCR T cells attract (solid lines, B-C) or both PyTCR and OT1 T cells attract (dashed lines, B-C) as well as data (bars) are shown. A simpler model in which only PyTCR T cells mediate attraction describes the data as well as the more complex model (likelihood ratio test, 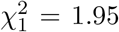, *p* = 0.16). Bars in panels B-C are the observed frequencies of parasites with a variable number of PyTCR (panel B) or OT1 (panel C) T cells. In panel D we show the data (points) and predictions of the model in which only Py-specific T cells attract all activated cells to the infection site (contours); model prediction is the log_10_ *P_ij_*(6) where *P_ij_* is the probability to observe *i* Py-specific and *j* nonspecific T cells around the parasite at *t* = 6h after T cell transfer. The point size represent the number of parasites having a given number of PyTCR/OT1 cells in 40 *μ*m radius, and thin dashed lines shows the line with slope = 1. In panel E points represent the prediction of the model on the number of PyTCR/OT1 T cells in clusters where *P_ij_* reaches maximum for *i* + *j* = const (see the contour plot in panel D). The solid line represents a regression line for the model predictions with slope = 0.76 which is significantly different from 1 (t-test, *p* < 0.001) and the dashed line shows the line with slope = 1. In panel F points represent co-clustering data of PyTCR and OT1 T cells around parasites (the same data are shown in panel D). Solid line represents a regression line with slope = 0.8 which is significantly different from 1 (t-test, *p* < 0.001). Results in panels D-F indicate model-predicted slight bias towards clustering of a larger number of PyTCR T cells which is not directly observed in the data (panel A).

Using the DD recruitment model we tested several different mechanisms of how specific and non-specific T cells may participate in cluster formation (see Table 1). Despite the highly correlated numbers of the Py-specific and non-specific T cells around Plasmodium liver stages (Figure 5A), different roles of these two CD8 T cell types seem to be inherent in the data (Table 2). Specifically, the model in which PyTCR T cells attract all cells into the cluster was statistically better at describing these data as compared to any other model tested (based on Akaike weights); interestingly, an alternative model in which OT1 cells exclusively drive cluster formation could not fit the co-clustering data well (model “Only OT1 cells recruit” in Table 2).

In two separate models we tested whether OT1 cells “help” in the formation of clusters which is driven by Py-specific T cells. Perhaps unsurprisingly in both models (“PyTCR and OT1 cells recruit at different rates” and “ PyTCR and OT1 cells recruit at different rates towards different cell types”) we found no evidence that OT1 cells enhance cluster formation (Table 2 and Figure 5B-C).

**Table 2:** Comparing alternative models which assume different contributions of Py-specific (PyTCR) and non-specific (OT1) T cells to cluster formation. We fit the basic mathematical model on co-clustering of Plasmodium-specific and non-speciific T cells (eqns. (12)–(15)) to the data on T cell clustering around Py liver stages assuming DD recruitment model and different mechanisms of how T cells contribute to cluster formation (see Table 1 for tested models). Here we list the estimated initial recruitment rate λ_0_ and how recruitment rate changes with cluster size λ_1_ (i.e., in the DD recruitment model the recruitment rate is λ*_k_* = λ_0_ + *kλ*_1_), the negative log-likelihood 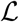, AIC, and Akaike weights w for the model fit. In these fits the total exit rate of T cells from the cluster of size *k* was fixed to *μ_k_* = 0.5*k*/h. In the column with estimates for λ_1_ we list specifically the predicted change in the cluster “attractiveness” by a given type of T cell (specific or non-specific) and towards a given type of T cells. For instance, an estimate λ_1_ = 0.58/h for the model in which only PyTCR cells recruit assumes that PyTCR cells recruit specific and nonspecific T cells at the same rate. In another model notation “PyTCR:OT1” denotes the recruitment rate induced by PyTCR cells for OT1 cells.

| Model | $\lambda_0$, 1/h | $\lambda_1$, 1/h | $\mathcal{L}$ | AIC | $w$ |
| --- | --- | --- | --- | --- | --- |
| Only PyTCR cells recruit | 0.14 | 0.58 | 128.0 | 260.3 | **0.40** |
| Only OT1 cells recruit | 0.15 | 0.60 | 131.4 | 267.8 | 0.01 |
| PyTCR and OT1 cells recruit at the same rate | 0.12 | 0.32 | 130.7 | 265.6 | 0.03 |
| PyTCR and OT1 cells recruit at different rates | 0.17 | PyTCR=0.67,<br>OT1=-0.16 | 127.1 | 260.6 | 0.34 |
| PyTCR and OT1 cells recruit at different rates towards different cell types | 0.18 | PyTCR:PyTCR= 0.73,<br>PyTCR:OT1= 0.61,<br>OT1:OT1= -0.15,<br>OT1:PyTCR= -0.23 | 126.7 | 264.8 | 0.04 |

In contrast, parameter estimates suggested that OT1 cells may inhibit cluster formation because the estimated OT1-driven recruitment rates λ_1_ were negative (Table 2); however, improvements of the fits of these two more complicated models were not supported by the likelihood ratio test (*p* > 0.1, see Figure 5B-C). Thus, our results suggest that non-specific T cells are “passive” participants in the clusters and do not significantly promote or impede the formation of clusters. A similar result was obtained recently using another Plasmodium experimental system [31].

Predictions of our best mathematical model in which only PyTCR cells recruit all activated T cells to the site of infection can be shown as the distribution of cluster sizes for each cell type (e.g., Figure 5B-C) as well as the probability to observe a cluster with *i* PyTCR and *j* OT1 cells (Figure 5D). Careful examination of this fit revealed that the model predicts a slight bias towards having more PyTCR cells per cluster than OT1 cells (Figure 5E). Linear regression analysis of the co-clustering data indeed suggests that there may be bias towards having more PyTCR cells than OT1 cells per cluster (Figure 5F); however, this result is not fully consistent with another analysis (e.g., Figure 5A), and the application of linear regression to data with integers may not fully appropriate. While the existence of such a bias is indeed in line with the analytical analysis of the steady state distribution of cluster sizes (see Supplement for mathematical proof), this bias is small (perhaps one extra PyTCR cell in clusters of a total size of 20), and biological relevance of such a bias for the killing of the parasite is most likely limited.

### 3.6 Clusters of Py-specific CD8 T cells around Py-infected hepatocytes are formed rapidly

Our analyses so far made an assumption that clusters around Plasmodium liver stages reach a steady state by 6-24 hours after T cell transfer. To understand potential limitations of this approach, we therefore performed several analyses using previously published experimental data.

Because our main mathematical model of T cell clustering (eqns. (1)–(2)) can be solved numerically and thus fitted to experimental data assuming a specific clustering mechanism (e.g., DD recruitment model), we investigated if the rates of T cell entry into the cluster (λ_0_ and λ_1_) and rates of exit from the cluster (*μ*) can be estimated from data in which PyTCR cell clusters around Py-infected hepatocytes were observed at 6 hours after T cell transfer. Interestingly, fitting the DD recruitment model (eqns. (1)–(2)) to data on the clustering of PyTCR cells transferred alone (Figure 2B or Figure 5A) revealed that model fits favored very high entry and exit rates, e.g., rates exceeding 20-30/h (results not shown). By fixing the exit rate from the cluster to multiple values we found that estimates of the absolute and relative values of the entry rate depended strongly on the exit rate values, and the relative entry rates (*θ*_0_ and *θ*_1_) approached constant values at high exit rates (Table 3). Importantly, all the fits of models with dramatically different exit rates were of nearly identical quality as based on negative log-likelihood suggesting that data on clustering of T cells at one time point are not sufficient to estimate entry and exit rates simultaneously. These results were confirmed for two independent datasets (experiments with PyTCR cells alone as shown in Figure 2A and Figure 5A) although exact values of parameter estimates such as *λ*_0_ did slightly vary between two sets of experiments (see Figure 2 and estimates in Table S1 in [29]).

**Table 3:**
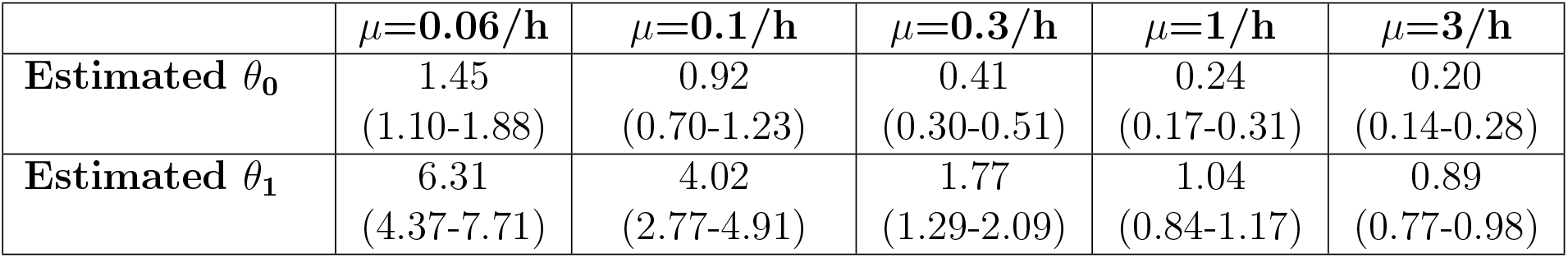
Estimated relative entry rates in the DD recruitment model (*θ*_0_ and *θ*_1_) strongly depend on the value of assumed exit rate from the cluster *μ*. We fixed the value of the rate of exit of T cells from the cluster *μ* to different values (indicated in the top row) and fitted the DD recruitment model (with λ*_k_* = λ_0_ + λ_1_*k* and *μ_k_* = *kμ*, see eqns. (1)–(2)) to experimental data on clustering of PyTCR T cells (*n* = 130) when transferred alone (cluster formation was observed 6 hours after T cell transfer, see Figure 5A). The quality of model fits to data as judged by the negative log-likelihood were nearly identical between different fits 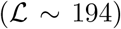; the fit of the model with *μ* = 3/h is shown in our previous publication (see Figure S1A and Table S1 in Cockburn *et al*. [29]). To compare parameter estimates we show relative entry rates *θ*_0_ = λ_0_/*μ* and *θ*_1_ = λ_1_/*μ*. Shown 95% confidence intervals for parameter estimates were obtained by bootstrapping the cluster data for individual parasites 1000 times. Interestingly, the ratio *θ*_1_/*θ*_0_ was relatively constant for different fits [48, and ms. in preparation].

| | $\mu=0.06/\text{h}$ | $\mu=0.1/\text{h}$ | $\mu=0.3/\text{h}$ | $\mu=1/\text{h}$ | $\mu=3/\text{h}$ |
| --- | --- | --- | --- | --- | --- |
| **Estimated $\theta_0$** | 1.45<br>(1.10-1.88) | 0.92<br>(0.70-1.23) | 0.41<br>(0.30-0.51) | 0.24<br>(0.17-0.31) | 0.20<br>(0.14-0.28) |
| **Estimated $\theta_1$** | 6.31<br>(4.37-7.71) | 4.02<br>(2.77-4.91) | 1.77<br>(1.29-2.09) | 1.04<br>(0.84-1.17) | 0.89<br>(0.77-0.98) |

To gain further insights into the kinetics of T cell cluster formation we analyzed additional data in which the same parasites (*n* = 32) were followed after T cell transfer over time and cluster sizes at different time points were recorded (Figure 6A and Figure S2). In these experiments imaging started between 4 and 8 hours after T cell transfer and followed for about 4 hours [29]. As expected there was a variable and statistically significant increase in the number of T cells found around individual Py-infected hepatocytes between T cell transfer and start of imaging (*t*_start_, Figure 6B). In contrast, in the following ~ 2 – 8 hours there was a minor change in cluster sizes (*t*_end_, Figure 6B). However, because imaging of CD8 T cell cluster formation started at different time points after T cell transfer there may be biases associated with the simple analysis of the data which takes into account only start and end time points of the clusters (e.g., Figure 6B). To obtained more accurate insights we further analyzed these data using mathematical models of cluster formation.

**Figure 6:**
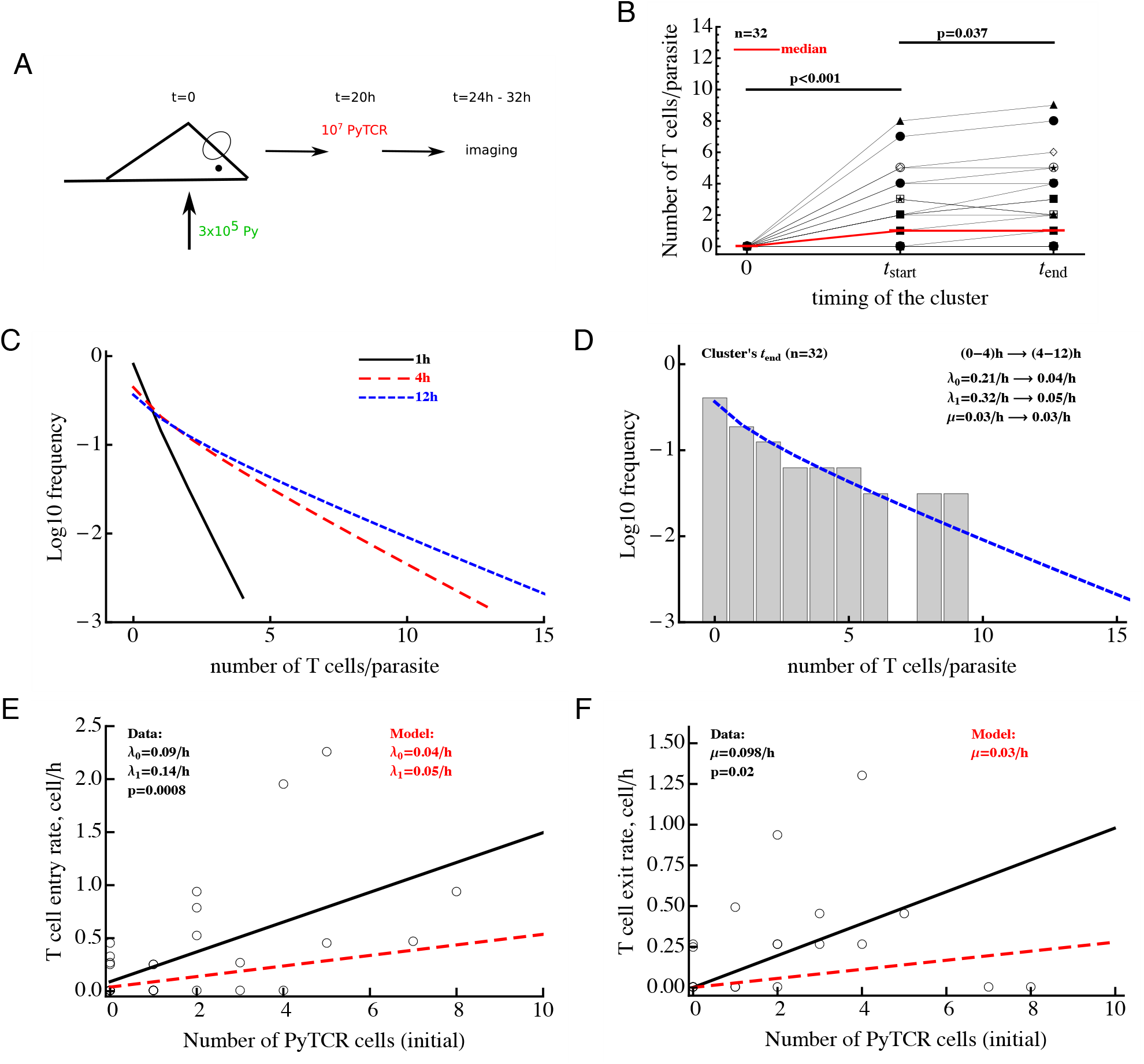
Clusters of T cells around the parasite are largely formed by 4 hours post T cell transfer. Panel A: mice were infected with 3 × 10^5^ GFP-expressing Py sporozoites. Twenty hours later 10^7^ Py-specific activated CD8 T cells (PyTCR) were transferred into infected mice and livers of these mice were imaged using intravital microscopy between 4 to 12 hours later. In total 32 parasites were randomly chosen and number of T cells in 40 *μ*m radius of the same parasites were counted at both times [29]. Panel B: significant increase in the median size of the cluster around Py-infected hepatocytes is observed in the first time period and there was a moderate increase in the median cluster size in the following 4-8 hours (Wilcoxon sum rank test). Thick red line shows change in the median number of T cells per parasite. In these experiments, 44% and 38% of all parasites did not have a single CD8 T cell nearby for first and last measurement of T cell clusters, respectively. Panel C: we plot the distribution of cluster sizes as predicted by the best fit model at different times after T cell cluster 4. The best fit model was a model assuming DD recruitment (eqns. (1)–(2)) with entry rates into the cluster being dependent on the time period (0-4h and 4-12h) but with the same exit rate during 12 hour period. Estimated parameters and their 95% confidence intervals for 0-4 h time interval are λ_0_ = 0.21 (0.11 – 0.34)/h and λ_1_ = 0.32 (0.11 – 0.49)/h; for 4-12h time interval are λ_0_ = 0.04 (0.0 – 0.10)/h and λ_1_ = 0.05 (0.02 – 0.08)/h with the exit rate *μ* = 0.030 (0.0 – 0.086)/h. Panel D: we show the observed distribution of cluster sizes at the last measurement for each parasite and predictions of the DD recruitment model for 12 hours after T cell transfer. Panels E&F: correlation between the T cell entry rate into the cluster (panel E) or exit rate from the cluster (panel F) as the function of the initial number of PyTCR T cells in the cluster. Points are experimentally measured values from Cockburn *et al*. [29], solid lines show the regression lines with estimated intercept λ_0_ = 0.09/h and slope λ_1_ = 0.14/h (panel E) or slope *μ* = 0.098/h (panel F); both slopes are significantly different from zero (t-test). Dashed lines in panels E&F show prediction of the mathematical model for the recruitment and exit rates estimated by fitting DD recruitment model to the clustering data.

To take full advantage of these “longitudinal” data in which T cell cluster formation was followed over time for individual parasites (Figure S2), we divided the data into individual “paths”, i.e., the number of T cells found near the parasite at sequential time points. For example, a parasite which did not have any T cells nearby and for which measurements were done at 0, 4 and 6.2 hours after T cell transfer, the path is “0 → 0 → 0”. For the parasite that was surrounded by most T cells in these experiments, the path is “0 → 8 → 9” for times 0, 4, and 6.2 h post T cell transfer (Figure 6B). A mathematical model of cluster formation can then be used to calculate the likelihood of a particular path by assuming that individual “sub-paths” along the path are independent (and thus by multiplying likelihoods of the model for individual sub-paths). For example, the probability to observe the path “0 → 8 → 9” at times (0, 4, 6.2) h is simply the product of the probability to observe 9 T cells in the cluster at 6.2 hours given that at 4 hours there were 8 T cells in the cluster and the probability to observe 8 T cells in the cluster at 4 hours given that at 0 hours there were 0 T cells in the cluster:

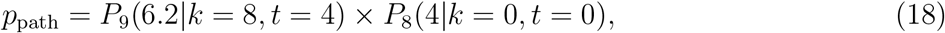

where the probability *P_k_*(*t*|*i*, *t*_0_) was calculated using the basic model (see eqns. (1)–(2)) with initial conditions *P_i_*(*t*_0_) = *θ_ij_* and *θ_ij_* is Kronicker delta (*θ_ij_* = 1 if *i* = *j* and *θ_ij_* = 0, otherwise). Fitting the DD recruitment model to these “longitudinal” data subdivided into “paths” resulted in the following entry/exit rates λ_0_ = 0.14/h, λ_1_ = 0.16/h, *μ* = 0.09/h. Additional analysis by fixing exit rate *μ* to different values and then comparing quality of the model fit using likelihood ratio test revealed that estimate of the parameters are relatively robust (i.e., fixing the exit rate to much lower or much higher values resulted in fits lower quality as judged by likelihood ratio test). Furthermore, by resampling the paths with replacement we found relatively small confidence intervals for the estimated parameters suggesting that measurement of T cell clusters longitudinally allows for a relatively accurate estimates of all three parameters of the DD recruitment model determining the kinetics of cluster formation (results not shown).

Parameter estimates of the model fitted to “longitudinal” (paths) data suggest that rates of entry into the cluster and exit from the cluster are relatively small, and this appears to contradict the formation of relatively large clusters already in 4 hours after T cell transfer (Figure 6C). Indeed, model fits did not accurately predict formation of clusters with > 5 T cells (results not shown). In addition, while the estimate of *θ*_0_ = λ_0_/*μ* was reasonable, the estimate of relative recruitment rate *θ*_1_ = λ_1_/*μ* was too low when compared with model estimates for clustering of T cells at 6 hours after transfer (e.g., Table 3 for *μ* = 0.1/h).

The major caveat of this analysis is the assumption that the parameters determining T cell clustering are constant over time. Our data indicate that formation of clusters may be slowing down over time (Figure 6B). Therefore, we fitted the DD recruitment model to the longitudinal/path data assuming that parameters determining kinetics of cluster formation depend on the time since T cell transfer. Given how the data were collected (Figure S2) for our analysis we made the simplest assumption that the rates are constant in two time intervals: (0 – 4) *h* and (4 – 12) *h* but may be different between the time intervals. Assuming that in the DD recruitment model recruitment rates λ_0_ and λ_1_ are time-dependent and the exit rate *μ* is time-independent, the model fitted the data significantly better than the DD recruitment model with constant parameters (likelihood ratio test, 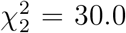, *p* < 10^-6^). Parameter estimates suggest a six fold reduction in both λ_0_ and λ_1_ 4 hours after T cell transfer (see legend of Figure 6 for actual parameter estimates). A similar decline in both λ_0_ and λ_1_ at 4 hours after T cell transfer was confirmed by fitting the model in which both rates declined by the same amount *α*; such a model fitted the data with a similar quality as the model that allowed for different decline in the two rates with time since T cell transfer (likelihood ratio test, 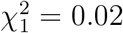, *p* = 0.89). Because the distribution of cluster sizes was measured experimentally at different time points it was not possible to visualize the model fits of the data. However, because model predictions suggested little change in cluster size distributions between 4 and 12 hours after T cell transfer (Figure 6C), the model predicted well the distribution of cluster sizes for each of the parasite at the end of imaging (Figure 6D, 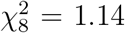, *p* = 1). Interestingly, this analysis indicated an extremely slow rate of T cell exit from clusters at 4-12 hours after T cell transfer suggesting that nearly every cell that enters the cluster after 4 hours post T cell transfer remains in the cluster which is an indirect support for the “retention” model.

An alternative DD recruitment model is one in which recruitment rates into the cluster remain constant over time but exit rates from the cluster change with time. This model did slightly improve the model fit of the data as compared to the model with constant parameters (likelihood change of 3.32, 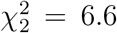, *p* = 0.01) and predicted that constant recruitment rates are λ_0_ = 0.14/h and λ_1_ = 0.19/h, and exit rate for the first 4 hours is *μ* = 0/h and for after 4 hours is *μ* = 0.19/h. This model suggests an alternative interpretation of the cluster formation dynamics — namely that T cells are recruited into the cluster and retained during the first four hours after T cell transfer — but after the initial time additional recruited T cells have a high chance of leaving. Because the quality of this model fit of the data was significantly worse than that of the model with time-dependent recruitment rates (ΔAIC = 21 or Akaike weight *w* < 0.001 for time-dependent exit rate model), our data appear to be more consistent with the time-dependent recruitment and constant exit. This suggests that the best explanation of the longitudinal clustering data is that the formation of clusters is driven by the DD recruitment model in which the rate of T cell recruitment into the cluster declines over time. Parameter estimates also suggest that the formation of clusters around Py-infected hepatocytes occurs mainly during the first 4 hours after T cell transfer.

The dynamics of change in the number of T cells near the parasite between 4 and 12 hours were followed by time-lapse intravital microscopy which allowed to calculate the number of T cells entering the cluster and leaving the cluster in this time period [29, Figure 6E-F]. Analysis showed that both entry and exit rates were strongly dependent on the cluster size k even though there was large variability in the number of T cells entering and exiting individual clusters. Interestingly, the slopes of the dependence of recruitment and exit rates was 2-3 fold higher for experimentally measured rates as compared to the parameters found by fitting DD recruitment model to longitudinal data (Figure 6E-F). One potential explanation of this difference is that perhaps not all cells that come near the parasite (i.e., within 40 *μ*m distance) recognize the infection and leave, thus, increasing the overall observed T cell surveillance rate. In contrast, the model only accounts for T cells which actually formed clusters and thus most likely have recognized the parasite.

## 4 Discussion

Studies from two independent groups showed that activated Plasmodium-specific CD8 T cells form clusters around Plasmodium-infected hepatocytes and that such clusters are correlated with elimination of the Plasmodium liver stages [29–31]. Application of mathematical models to data on distribution of the number of Py-specific CD8 T cells around randomly chosen parasites suggested that formation of the clusters is not a random process; the model in which activated T cells of different specificities are attracted at a rate proportional to the number of Py-specific T cells already present near the parasite, described the data with best quality [29]. More recent work also suggested that formation of CD8 T cell clusters around Plasmodium-infected hepatocytes depends on CD11c^+^ cells and that activated CD8 T cells, specific to irrelevant antigens, do not appear to play a significant role in protection against Plasmodium challenge [31]. We analyzed experimental data with the use of mathematical modeling to provide further insights into potential mechanisms of the formation of clusters around Py-infected hepatocytes.

First, we found that several independent experimental datasets are fully consistent with the model in which variability in the number of activated Py-specific CD8 T cells located near the parasite-infected hepatocytes is driven by variability in the “environment” around the infected hepatocytes, providing indirect support for the T cell-extrinsic mechanism of cluster formation (e.g., Figure 3). These results suggested that data on clustering of Py-specific T cells alone may be insufficient to discriminate between T cell-intrinsic and T cell-extrinsic mechanisms of cluster formation [49]. A key experiment, rejecting the “variability of the environment” hypothesis as the sufficient mechanism explaining distribution of cluster sizes is one involving either transfer of only OT1 T cells (which are not specific to Py antigens) or OT1 T cells together with PyTCR T cells – only in the latter case, OT1 T cells form co-clusters with Py-specific T cells (Figure 4 and see [29, 31]). The mathematical model assuming fixed yet variable (between individual parasites) environment was not able to accurately explain such data (Figure 4). The result, however, does not mean that inflammation is irrelevant for parasite’s replication in the liver. In fact, recent work suggested that sporozoite infection of the liver does lead to inflammation [36].

Second, while OT1 T cells of irrelevant specificity are found in clusters together with Py-specific CD8 T cells, we found no evidence that OT1 improve cluster formation (Figure 5). If anything, OT1 T cells may in fact reduce the rate of recruitment of other T cells into the cluster as indicated by the negative values for the recruitment rate λ_1_ (Table 2); however, this value was not significantly different from zero. Mathematical modeling also suggested that there may be a slight bias in the clusters to have more Py-specific T cells than T cells of irrelevant specificity per cluster but the biological relevance of such a small bias remains unclear. The limited role of T cells of irrelevant specificity in the formation of T cell clusters in Py-infected mice is consistent with the observation that transfers of large numbers of activated CD8 T cells with irrelevant specificity into Plasmodium-infected mice did not impact efficiency at which parasites were killed by Plasmodium-specific T cells [31]. Interestingly, and perhaps surprisingly, this result contradicts a recent observation of suppression of development of T-cell-driven type 1 diabetes by islet-non specific CD8 T cells [50].

Third, by following longitudinal changes in the number of CD8 T cells around individual parasites over time we found that T cell clusters are formed rather rapidly, at least within the first 4 hours after T cell transfer (Figure 6B), and mathematical modeling predicted recruitment of T cells to the parasite and retaining of the T cells in the cluster in that time period. Interestingly, the rates of entry into and exit from the clusters declined after the 4 hours six fold further supporting the conclusion that clusters are formed rapidly and few cells enter and exit the cluster after 4 hours since T cell transfer (Figure 6C-D). Stochastic simulations of the formation of clusters assuming DD recruitment model with different entry/exit rates also suggested that between 4 and 8 hours post T cell transfer, entry and exit rates cannot be large (Figure S3). This is because when these rates are large, changes in the cluster size in the 4-8 hour time period are highly variable with some clusters growing in size exponentially while other clusters nearly disappearing (e.g., Figure S3C&F). This, however, was not observed in experimental data (shown by dotted histogram in Figure S3D-F and see Figure 6B and Figure S2). Rapid recruitment of CD8 T cells to the liver stages in the first 4 hours after T cell transfer may be the result of the specific experimental set-up as it is expected that immediately after intravenous injection, large numbers of T cells would be passing through the liver increasing chances of T cells finding the infection site [51], and that the number of liver-resident CD8 T cells tends to reach a steady state at 2-3 hours after T cell transfer [52, James O’Connor and Ian Cockburn (ms. in preparation)].

Interestingly, there was some discrepancy between the estimated rates of T cell entry into the cluster and exit from the cluster measured experimentally and predicted by the model (Figure 6E-F) most likely indicating that not all T cells that were observed to come in close proximity with the parasite recognize it. We also found a strong correlation between experimentally measured rates of T cell entry into and exit from the clusters (Figure S4) which may indicate that in addition to T cell-intrinsic mechanisms of clustering, some parasites may be more “attractive” to T cells. Indeed, none of our tested models could well explain the formation of extremely large T cell clusters around Py-infected hepatocytes (e.g., with 15 or more T cells, see Figure 2B) which could indicate the need for future models to include both DD recruitment and variability in parasite’s attractiveness.

In this paper we analyzed a number of different datasets that involve different cell types, different experimental set-ups, and different mice. We found it encouraging that some of these datasets were in a way “consistent”. Specifically, we observed similar clustering of CD8 T cells in naive mice (Figure 3A), PT-treated PyTCR T cells (Figure 2B), or activated OT1 T cells of irrelevant specificity (Figure 4A) and the random entry/exit model described these data with nearly identical parameters (likelihood ratio test, 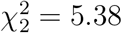, *p* = 0.07). The DD model could describe the distribution of cluster sizes of PyTCR T cells in three different experiments (Figure 2A and data in [29]) with identical parameters (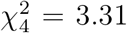, *p* = 0.51). However, the clustering of CD8 T cells following immunization with radiation-attenuated sporozoites (Figure 3B) did not match well the clustering of the mixture of PyTCR and OT1 T cells (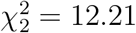, *p* = 0.002) perhaps highlighting potential differences between active and passive immunizations (the latter involving transfer of pre-activated CD8 T cells).

In multiple analyses we found that a DI exit (retention) model did not describe well the clustering data. However, a poorer fit of the model (as compared to other tested models) does not necessarily falsify a model [49], and additional experiments will be needed to formally rule out this model. Fitting the DI exit model to the “longitudinal” data on change in cluster size around individual parasites (e.g., Figure S2) revealed that this model could not accurately describe the data assuming a constant entry and time-dependent exit rates based on likelihood of the model (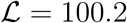 vs. 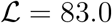 of the DD recruitment model with time-dependent recruitment and constant exit rates, results not shown). In addition, if the rate of T cell exit from the clusters found in the DD recruitment model is constant over the course of the first 12 hours since T cell transfer, it would suggest that T cells mostly enter the clusters and rarely exit them (given *μ* = 0.028/h corresponding to the residence time of T cells in the cluster of about 1/*μ* ≈ 36 h), providing some indirect support for the retention model.

Conversely, our result that the DD recruitment model describes most of the data with best quality does not prove that this model is the true mechanism of the formation of large clusters around Py-infected hepatocytes. Future experiments will have to test the major prediction of the model — that clusters of a large size attract more T cells per unit of time. Such experiments may involve measurement of T cell movements in the liver using intravital microscopy and estimating bias in T cell shift towards the parasite. Indeed, our recent work suggested that there is a bias in PyTCR cell movements towards Py-infected cells [34] but more analyses are needed to evaluate whether such a bias depends on the number of T cells already present at the parasite and whether a small bias is sufficient to explain the formation of larger clusters (with *k* ~ 5 – 10 of CD8 T cells per parasite) within few hours after T cell transfer. Detecting a bias in T cell movement towards the infection site may be complicated as our current analysis predicts that “attraction” seems to be present only during the first 4 hours after T cell transfer (Figure 6C-D).

Our analysis has several potential limitations. The biggest issue is that by using numerical solutions of the DD recruitment model we showed that the distribution of cluster sizes at a single time point does not allow to accurately estimate the rates of T cell entry into the clusters and T cell exit from the clusters, and thus, most of our analysis were restricted to estimating relative recruitment rates. Ongoing analysis based on analytical solutions in Bailey [48] has also demonstrated this point using analytical derivations (ms. in preparation). While the estimated values of the recruitment rates λ_0_ and λ_1_ in the DD recruitment model directly depend on the assumed exit rate *μ* (see Table 3) we showed that the likelihood of the model fit to data assuming a steady state or dynamics for clusters at a given exit rate *μ* were nearly identical strongly suggesting that our results on best fit models obtained assuming steady states are robust. However, the actual values of the entry and exit rates cannot be found with certainty as these depend on the actual value of the assumed exit rate (Table 3).

Another complexity in the analysis comes from our finding that rates of T cell entry into the cluster are time-dependent (Figure 6). To investigate whether this impacts our selection of best fit models assuming steady state solutions we did the following. We fitted the DD recruitment model to the clustering data at one time point by assuming that early recruitment rates λ_0_ and λ_1_ are unknown and that late entry rates are fixed to values found from the analysis of longitudinal data (Figure 6) and that the exit rate *μ* is constant. Under these minimal assumptions the model fit was of nearly identical quality as the model fit of the data assuming a steady state (results not shown). Therefore, even for time-dependent parameters our results determining which models are not consistent with clustering data remain valid.

An important experimental limitation of our data is the way of how experiments were performed whereby pre-activated T cells were transferred into mice that had been already infected with Plasmodium sporozoites (e.g., Figure 2A). This sequence of events does not fully match the physiological situation in which activated or memory CD8 T cells are already present at the time of sporozoite challenge. In fact, the rapid predicted decline in the rates of T cell recruitment into clusters with time suggests that it may be an artifact of the experimental system. Whether change in the experimental protocol will lead to support of the same mathematical models of cluster formation remains to be determined (and is the focus of our ongoing experiments and analyses).

Mathematical methodologies used in this work provided deeper understanding of how CD8 T cells form clusters around Plasmodium-infected hepatocytes. While formation of such clusters was a novel observation in malaria infection of the liver, clusters of immune cells have been observed in multiple systems including herpes simplex virus (HSV) [53] and Mycrobacterium tuberculosis (Mtb) [54]. In fact, formation of granulomas in Mtb-infected animals and humans is a classical example of T cell clustering around the infection site. Interestingly, both Mtb-specific CD4 T cells and CD4 T cells of irrelevant specificities were found in granulomas of Mtb-infected monkeys [55] which could be explained by the DD recruitment model extended in this work. Movement of neutrophils towards an injury site may also depend on the number of neutrophils that have already reached the site [56]. It may be useful to combine mathematical modeling tools for deeper understanding of the mechanisms of formation of clusters of immune cells in these and other systems.

While our work provides some clarification regarding mechanisms of CD8 T cell cluster formation around Plasmodium-infected hepatocytes, many questions remain. In particular, while clusters appear to be important for the death of the parasite [29, 30], whether clusters of a larger size kill the parasites faster remains unknown. Classical work involving killing of chromium-labeled target cells by cytotoxic T lymphocytes (CTLs) suggested a faster killing of targets bound by multiple CTLs [57], and *in vivo*, death of peptide-pulsed targets is directly proportional to the number of peptide-specific CTLs [33]. Recent work also suggested that the probability of death of virus-infected cells in skin in vivo was higher when the infected cell was contacted by several antigen-specific CD8 T cells [32]. Whether the same relationship holds for T cells killing Plasmodium parasites in the liver remains to be determined. However, several studies have shown that probability of clearance of i.v. injected Plasmodium sporozoites does depend on the number of vaccine-induced CD8 T cells in the liver [22, 23, 58]. However, these previous studies are numerically inconsistent suggesting that either 3 × 10^4^ [22] or 10^6^ [23] memory CD8 T cells in the liver are needed for sterilizing protection. Further work is required to accurately quantify the number of T cells needed for protection.

Our results suggest that activated CD8 T cells of irrelevant specificities do not play a major role in cluster formation, and elegant experiments demonstrated that large numbers of non-specific T cells do not impair the ability of Plasmodium-specific T cells to eliminate the parasites [31]. However, the latter result was found by using only two different ratios of Plasmodium-specific and non-specific T cells and 3 mice per group, so it remains to be determined if competition between such cells for the access to infected cells occurs at higher ratios, e.g., as has been observed in another system [50]. In natural settings we do expect that Plasmodium-specific T cells will be likely outnumbered by memory T cells specific to other infections, and therefore, deeper understanding of such competition may be of relevance to malaria vaccines, inducing liver-resident memory CD8 T cells for protection [59].

Accumulation of large numbers of CD8 T cells around Plasmodium-infected cells raises an intriguing possibility that parasites may in fact attempt to attract T cells. While this may be detrimental to an individual parasite, as a population this may give an advantage if attracting many T cells to one site prevents T cells from effectively locating parasites at other sites. To cause blood-stage infection, there is a need for only one liver stage to mature and release differentiated parasites (merozoites) into the blood stream. Indeed, it should be noted that in most of our experiments, many of surveyed parasites did not have a T cell nearby at 6-8 hours after T cell transfer. Future studies may be needed to investigate whether such a strategy is indeed evolutionarily advantageous.

It remains unclear how relevant our results are for T cell-mediated protection of humans against malaria. Because of the need of imaging i.v. injected sporozoites in the liver, large numbers of parasites must be used. This is in contrast with very few sporozoites that humans are likely to be exposed to when bit by infectious mosquitoes.

Taken together, here we illustrated the power of combining the use of detailed quantitative experimental data with mathematical modeling, and limitations that come from inability to make solid conclusions from extensive yet limited experimental data. The field of immunology will likely benefit from closer collaborations between experimentalists and modellers where experimentalists being involved in data analyses and modeling, and modellers are cooperating with experimentalists in designing experiments to test and potentially falsify alternative mathematical models.

## 5 Acknowledgements

This work was supported by the NIH grant (R01 GM118553) to VVG. This manuscript has been released as a preprint at BioArxiv [60].

## 6 Author contributions

RKK, IAC, and VVG designed the study. RKK and VVG run most of analyses and wrote first drafts of the paper. HR confirmed some of the numerical results and provided derivation of the bias in co-clustering of antigen-specific and non-specific T cells. IAC performed additional data analyses and provided feedback on data interpretation and modeling. The final version of the paper was written by VVG but all authors contributed to the writing of the final version of the paper via comments and suggestions.

## Supplemental Information

### Bias in clustering of Plasmodium-specific CD8 T cells and T cells of irrelevant specificity

In fitting mathematical models to data on co-clustering of PyTCR (Py-specific) and OT1 (OVA-specific) CD8 T cells we found slight bias in the number of PyTCR cells in a given cluster. Here we provide mathematical justification of this observation.

#### Deflning the problem

The linear system of ODEs given in eqns. (1)–(2) can be written as follows:

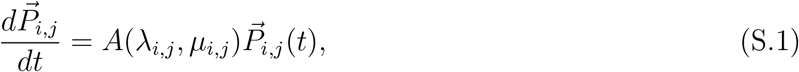

where *A*(λ_*i,j*_, *μ_i,j_*) is the probability transition matrix of time-independent entry and exit rate parameters *λ_i,j_* and *μ_i,j_*, respectively, and the subscript (*i, j*) denotes the number of PyTCR and OT1 cells of a given cluster, respectively, specific to the probabilities and rate parameters. Here, we consider the PyTCR density-dependent recruitment co-cluster model, where we assume that entry rates λ_*i,j*_ of cell combinations (*i, j*) are time-constant, and both PyTCR and OT1 cells are attracted to a cluster depending on the density of PyTCR cells (type *i*) in the cluster, which is called the DD recruitment model. Thus, we can write λ_*i,j*_ = λ_0_ + *i*λ_1_; where λ_0_ and λ_1_ are parameters. The exit rate μ_*i,j*_ from a cluster are per-capita with respect to the particular cell density, i.e., *μ_i,j_* = *iμ* or *μ_i,j_* = *jμ*, where *μ* > 0 is a constant.

Defining *θ*_0_ = λ_0_/*μ, θ*_1_ = λ_1_/*μ*, we can write

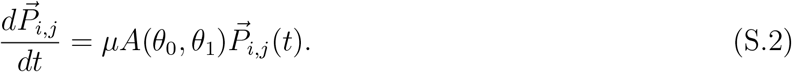

Consider the solutions 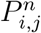 of the above system of equations written in a matrix of size 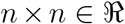, with the largest possible co-cluster size *k_max_* = n, i.e., max(*i + j*) = *n*, where *i* (i.e., PyTCR cells in a cluster) denotes the rows, and *j* (i.e., OT1 cells in a cluster) denotes the columns of the matrix. The previous results of fitting DD recruitment model to the data showed that probabilities *P_i,j_* of entries (*i, j*) for smaller *j* of the upper right triangle of the matrix are likely to be larger than their respective mirror entries (*j, i*) of the lower left triangle of the matrix, and in contrast, *P_i.j_* of entries (*i, j*) for larger *j* of the upper right triangle of the matrix are likely to be larger than their respective mirror entries (*j, i*) of the lower left triangle of the respective matrix. That is, the probabilities for smaller clusters to have a larger number of OT1 cells are greater, whereas the probabilities for larger clusters to have a larger number of PyTCR cells are greater. Here, we show the formation of this systematic bias in the DD recruitment model for *P_i,j_* solutions for both the steady state and time-evolving conditions.

#### Explanation of bias for the system in steady state

Consider the system at steady state, i. e., where the rate of evolution of probabilities of all possible combinations of PyTCR and OT1 cells over time in the ODEs are zero. Thus, we get *A*(*θ*_0_,*θ*_1_)*P_i,j_*(*t*) = 0 for *μ* > 0. Note below that superscript *n* of 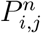 indicates the maximum cluster size of the matrix.

**I) Case** *n* =1 [i.e., max(*i* + *j*) = 1].

**Table S1:**
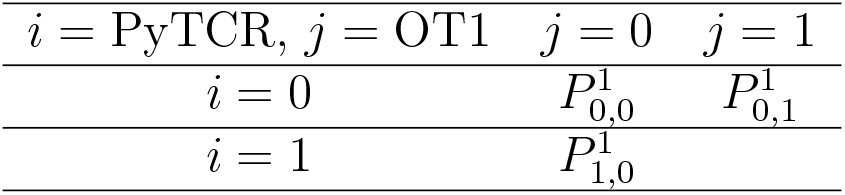
Probability table for *n* = 1.

| $i = \text{PyTCR}, j = \text{OT1}$ | $j = 0$ | $j = 1$ |
| --- | --- | --- |
| $i = 0$ | $P_{0,0}^1$ | $P_{0,1}^1$ |
| $i = 1$ | $P_{1,0}^1$ | |

The steady state equations of the *n* = 2 matrix system are

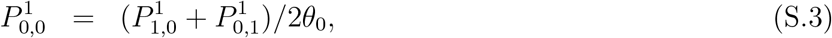

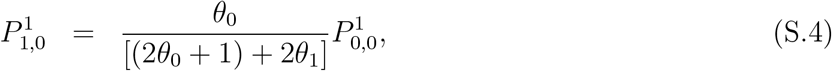

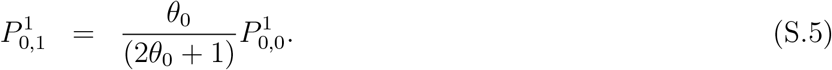

Since *θ*_0_, *θ*_1_ > 0, we get 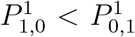. Here, 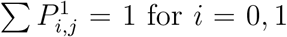 and *j* = 0,1, s.t. (*i + j*) ≤ 1. And, the ratio 2*θ*^1^/*θ*^0^ determines how large the 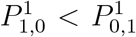 relationship is. Also, note that for any system of equations of size max(*i + j*) = *n* at steady state, the relationship in eq. (S.3) satisfies for probabilities 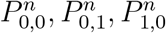. That is,

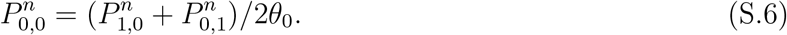

##### Corollary I

Due to similarity of coeffcients between the linear equations (S.3) and (S.6) of a given system, regardless of the matrix size n, s.t., max(*i + j*) = *n*, with fixed rate parameters, 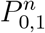 and 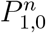 given by eq. ((S.6)) can be expressed as 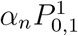 and 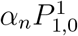, respectively, for any given 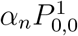 of the eq. (S.3), where 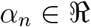. Here, 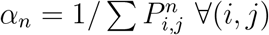 entries.

Thus, the condition 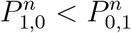 holds true for every solution of a given system of size *n*.

**II) Case** *n* = 2 [i.e., max(*i + j*) = 2].

Here, we get,

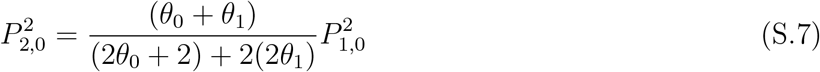

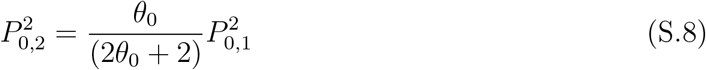

Since we can rewrite 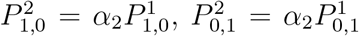 and 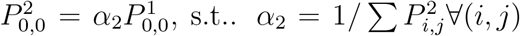 entries, s.t., max(*i + j*) = 2, from Corollary I, we get

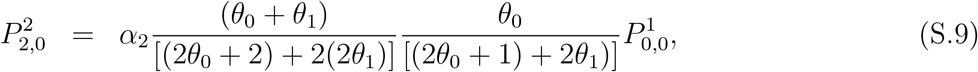

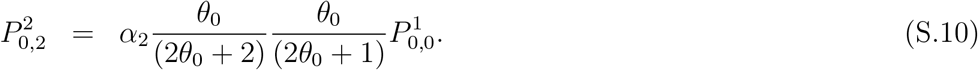

**Table S2:** Probability table for *n* = 2.

| $i = \text{PyTCR}, j = \text{OT1}$ | $j = 0$ | $j = 1$ | $j = 2$ |
| --- | --- | --- | --- |
| $i = 0$ | $P_{0,0}^2$ | $P_{0,1}^2$ | $P_{0,2}^2$ |
| $i = 1$ | $P_{1,0}^2$ | $P_{1,1}^2$ | |
| $i = 2$ | $P_{2,0}^2$ | | |

##### Corollary II

If (*x < y*) and (*α* < *β*) then (*x* + *α*)/(*y* + *β*) > (*x/y*) only if (*x/y* < *α/β*)

It follows from Corollary II that in eqns. (S.9) and (S.10), 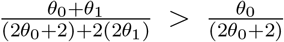, and also, 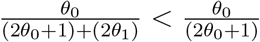. Thus, whether 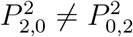 is determined by the condition 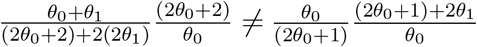, which simplifies to 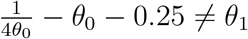. Here *θ*_0_ < 1 in general.

##### Corollary III

It follows from Corollary I, in general, that for a given system of matrix size *n* = *h* +1, i.e., max(*i + j*) = *h* +1, the 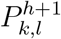 for max(*i + j*) = *h* – 1 are given as functions of entries 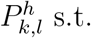 s.t., max(*k + l*) = *h*, multiplied by the factor *α*_*h*+1_, where 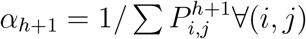.

Thus, for the general case for matrix max(*i + j*) = *h*, comparing 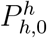 vs. 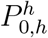, we get

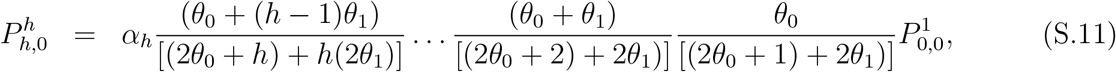

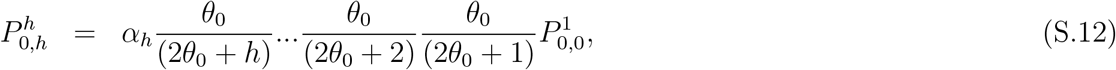

for constant *α_h_*. Note that equations (S.11) and (S.12) are functions of 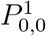.

**Table S3:** Probability table for *n* = *h*.

| $i = \text{PyTCR}, j = \text{OT1}$ | $j = 0$ | $j = 1$ | $j = 2$ | … | $j = h - 1$ | $j = h$ |
| --- | --- | --- | --- | --- | --- | --- |
| $i = 0$ | $P_{0,0}^h$ | $P_{0,1}^h$ | $P_{0,2}^h$ | … | $P_{0,h-1}^h$ | $P_{0,h}^h$ |
| $i = 1$ | $P_{1,0}^h$ | $P_{1,1}^h$ | $P_{1,2}^h$ | … | $P_{0,h-1}^h$ | |
| ⋮ | ⋮ | ⋮ | $\ddots$ | … | | |
| $i = h - 1$ | $P_{h-1,0}^h$ | $P_{h-1,1}^h$ | | | | |
| $i = h$ | $P_{h,0}^h$ | | | | | |

And, thus the 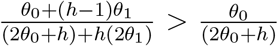 holds true depending on the condition if

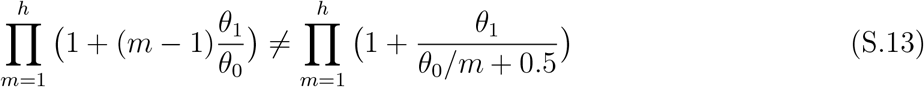

as every multiplicative terms, factors after *h* > 2 has a potential to yield 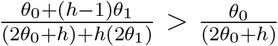 for large h, following the Corollary II, for 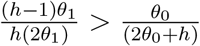, i.e., 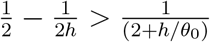 for 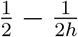 goes from 0 to 1/2, while 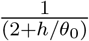 goes from 1/2 to zero for h tending from 1 to infinite. In other words, when *h* gets larger (*h* ≫ 1), L.H.S of condition in equation (S.13) turns greater than R.H.S of the condition.

It follows that for small clusters with only one type of cells, the probabilities for the number of OT1 cells to be greater than that of PyTCR cells is greater, i.e., 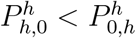, whereas for large clusters the probabilities for the number of PyTCR cells to be greater than that of OT1 cells is greater, i.e., 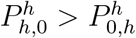.

For the case of diagonal entries of matrix size max(*i* + *j*) = 3, we can write

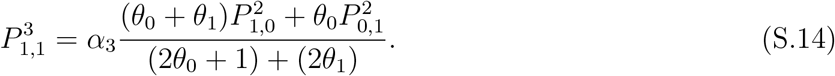

Here, the proportional contribution of 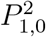 on 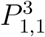 is greater than that by 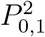 on 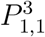, whereas, we also get that the proportion of exits from 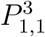 towards 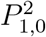 is less than that of 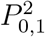 as 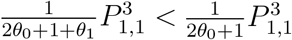, from the respective functions of 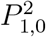 and 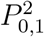 from steady state equations. Therefore, there is a net differential (flow) of probabilities from PyTCR cells towards the cells with more OT1 cells of the matrix size *n* = 3.

Furthermore, for the general case of the above, i.e., diagonal entries of matrix size max(*i* + *j*) = 2*h* + 1 (odd number), we can show that

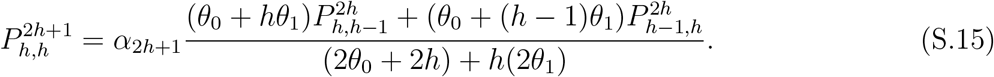

Here, when *h* → ∞, the proportional contribution from the entries of 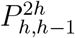 and 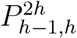 of matrix *n* = 2*h* converges to a single value (constant). The convergence is also true for the proportionate exits from 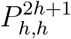 into the two entries 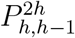 and 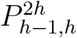. Thus, it follows that when h is larger, the flow of probability from PyTCR cells towards the cells with more OT1 cells converges to zero.

Any probabilities of mirror entries on either side of (*h, h*) entry of large *h*, we note that

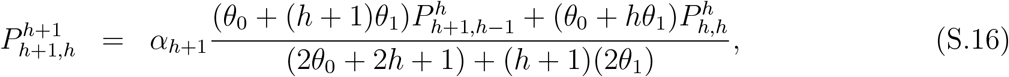

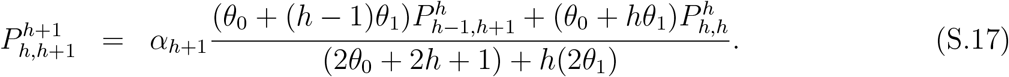

As before, as *h* increases, the contribution of 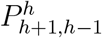 towards 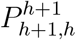 converges to that of 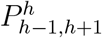 towards 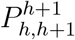 as per the Corollary II. Thus, for large *h*, 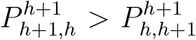 depending on 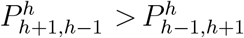, for contributions of 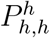 towards 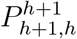 and 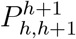 converge.

Furthermore, the special case of any two mirror entries (*h*, 1) and (1, *h*) of matrix size max(*i* + *j*) = *h* +1

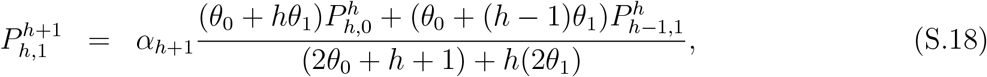

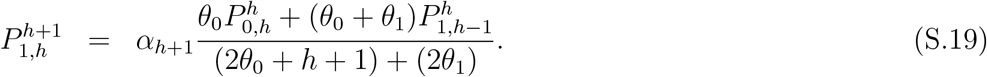

By Corollary II, here the coeffcients of respective entries in function 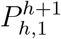 are greater than that in the function of 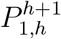 for larger *h*. Thus, it follows that 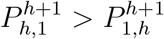 as 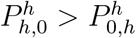 and 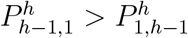 as *h* gets larger.

For any two mirror entries (*i, j*) and (*j, i*) for the case max(*i* + *j*) = 3 matrix, we can write

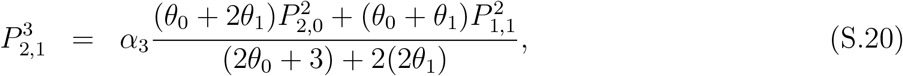

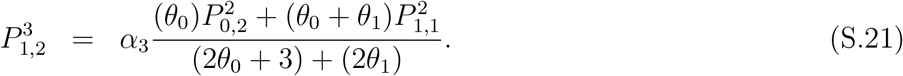

By comparing coefficients of equations (S.20) and (S.21), we note that the contribution of 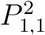 towards 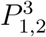 is greater than that towards 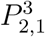. From the same, the contribution of 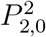 towards 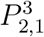 is greater than that of 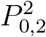 towards 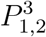 as per the Corollary II. Thus, if 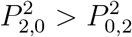, then 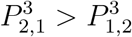, depending on the magnitude of the contribution by 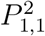, towards each, and vice-versa.

For the general case of the above special case, for any (*k, 1*) and (*l, k*) entries, s.t. (*l* + *k*) = *h*, and *l* < *k* we get

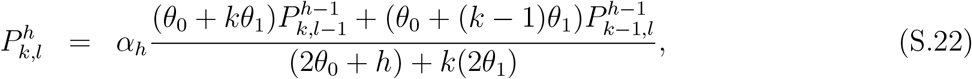

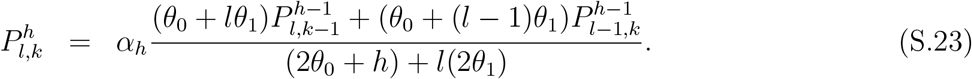

For *k* > *l*, using Corollary II, we get that the coefficients of respective entries in 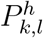 are greater than those of mirror entries in 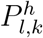. Thus, it follows that 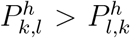 if 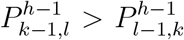 and 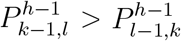 as *h* gets larger (*h* ≫ 1), also because the flow of probability through 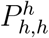 tends to zero for large *h* as we have shown above.

Thus, once the probability entries in the lower left triangle, i.e., where PyTCR cell numbers are higher, turn greater than those mirror entries in the upper right triangle, i.e., where OT1 cell numbers are higher, depending on the condition in equation (S.13), and those follow similarly, the pattern of bias remains the same for large *h* regardless of how large the matrix is.

#### Explanation of the bias for the system of evolving probabilities over time

The solution to the system at time *t*_2_ = *t*_1_ + *δt*, evolved from time *t*_1_ yield 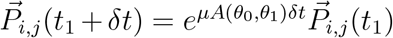 where *e* denotes the matrix exponent with elements ∈ ℜ, can be written as 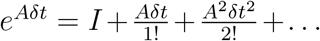, where I is the identity matrix. Assuming higher orders (> 1) of the series expansion are negligible compared to its first order solution, we can write the solution to the system, evolved from time *t*_1_ to *t*_2_ as 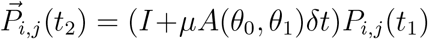, for elements in A and t are < 1. This yields 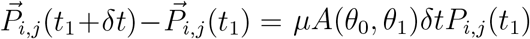. Note that the steady state solution was obtained by setting this difference to 0. We can write the above as 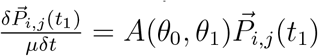.

Thus, the solutions of 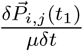 of the above to each cluster combination (*i, j*) as a function of 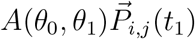, can be written as, for example from equation (S.3), for the case *n* = 1, 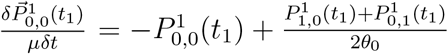, and similarly from equation 2, 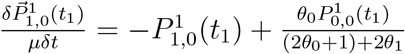, and so on ∀(*i, j*) combinations given for the case *n* = 1 and so forth for all *n* = 2, 3, 4,…. These can be also written as 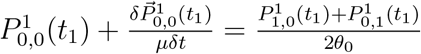, and 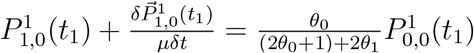 and so forth for all *n* = 2, 3, 4,…. Therefore, the patterns in bias, explained under the steady state solutions, 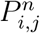 in the main text are qualitatively similar for solutions 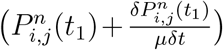 for all combinations of (*i, j*), and also for matrices *A* of any size *n*. This is because 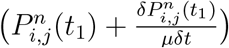 are functionally equivalent to 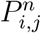 solutions under the steady sate condition. Thus, the bias discussed in the main text is similar for any 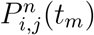 computed discretely in the form of 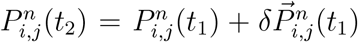 step-wise, starting from *t* = 0 to any *t*, choosing *δt*, s.t., *μδt* = 1.

**Figure S1.**
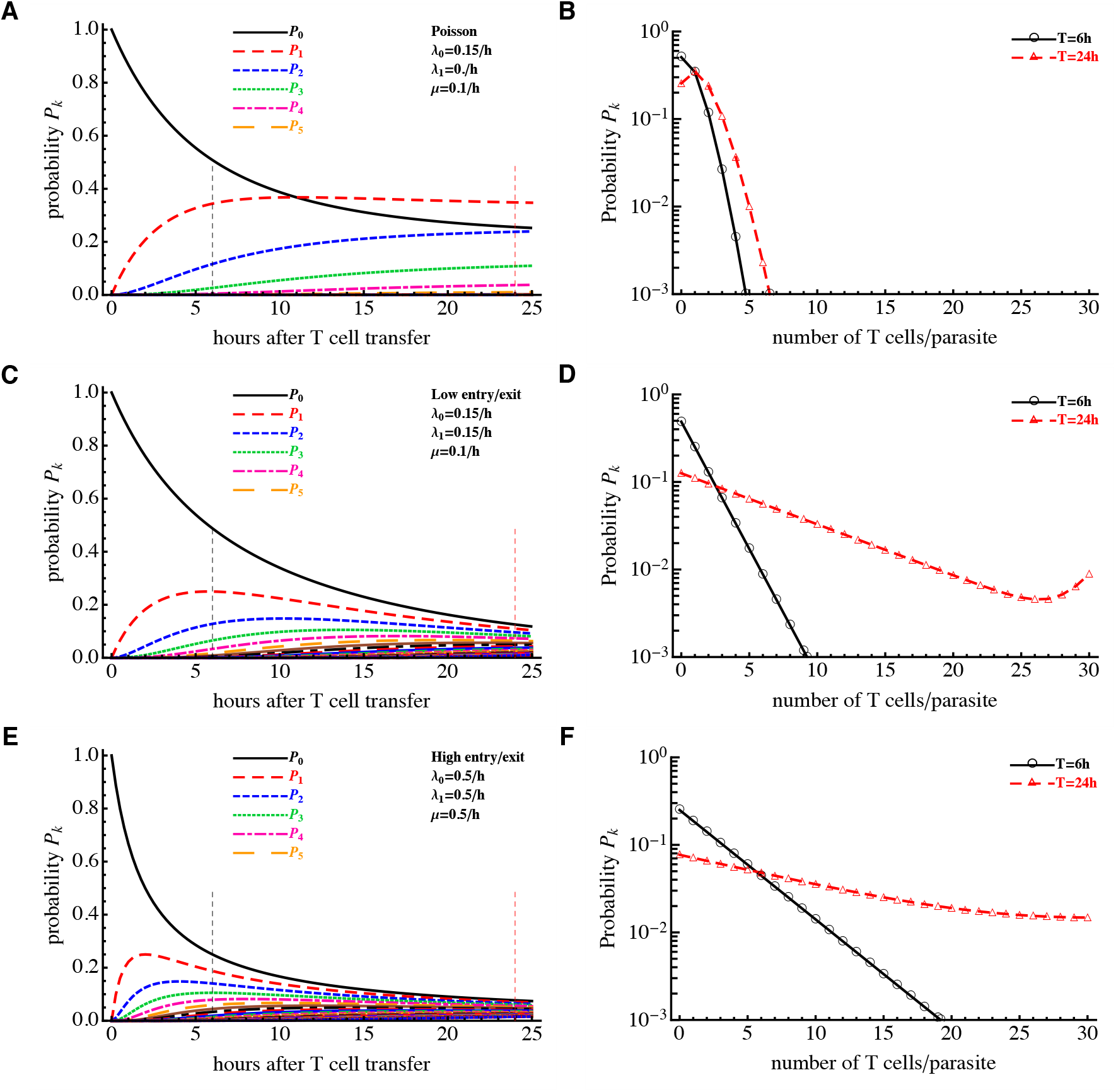
Dynamics of cluster formation as predicted by the random entry/exit (Poisson) and density-dependent (DD) recruitment models. We simulated the dynamics of cluster formation using eqns. (1)–(2) in the main text and plot the probability to observe *k* T cells around a parasite, *P_k_*(*t*), as predicted by the Poisson model (panels A-B, λ_0_ = 0.15/h, *μ* = 0.1/h), DD recruitment model with low entry into/exit from the cluster rats (panels C-D, λ_0_ = 0.15/h, λ_1_ = 0.15/h, *μ* = 0.1/h), or the DD recruitment model with high entry into/exit from the cluster rates (panels E-F, λ_0_ = 0.5/h, λ_1_ = 0.5/h, *μ* = 0.5/h). We show the dynamics of the probability *P_k_* over time (panels A, C, E) or the distribution of cluster sizes at 6 or 24 hours after start of cluster formation (panels B, D, F).

**Figure S2.**
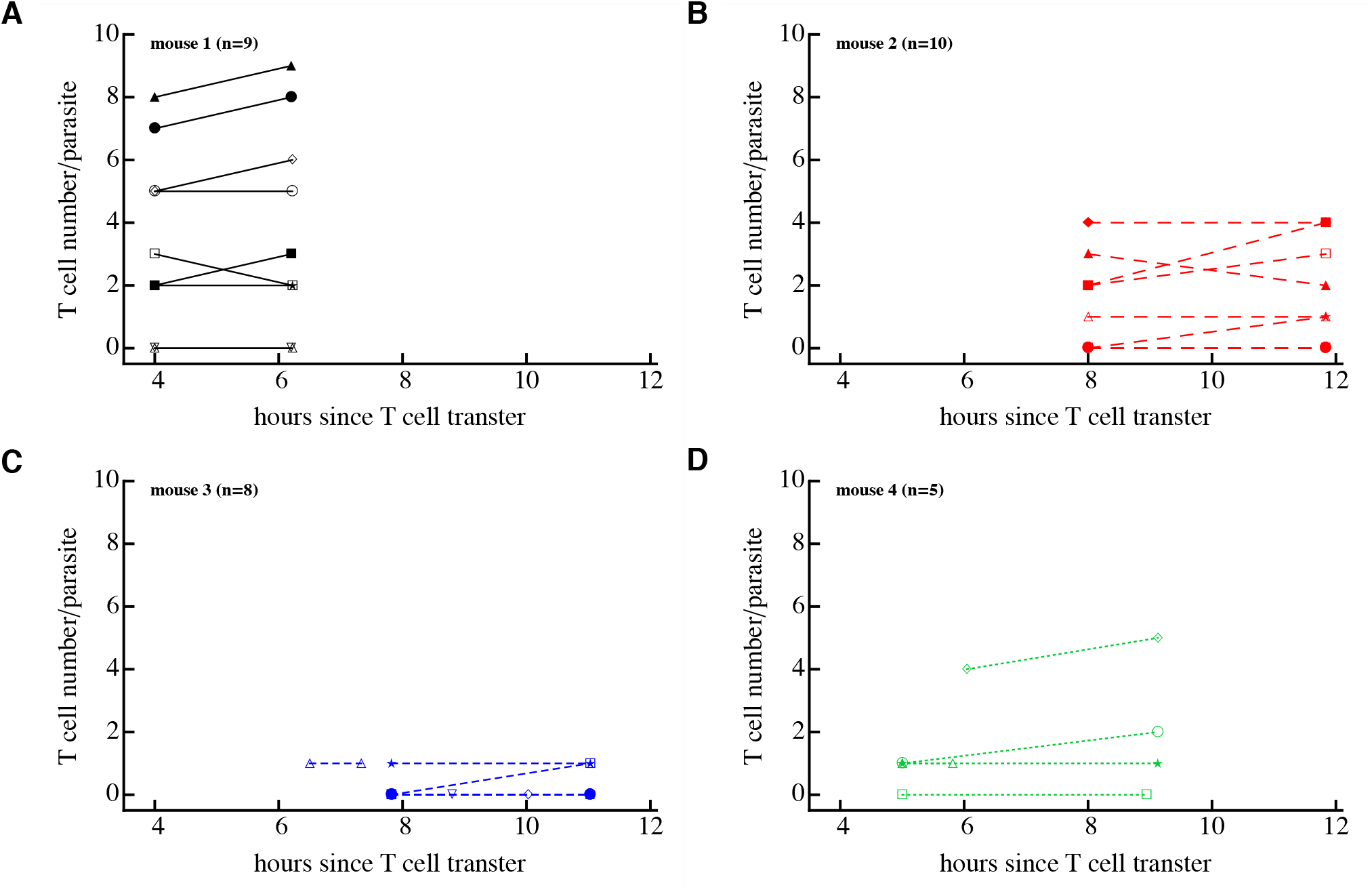
Moderate change in the T cell cluster size over time. We performed imaging experiments as described in Figure 6A and counted the number of T cells found around individual parasites at start and end of intravital imaging done after T cell transfer. Individual panels show change in T cell cluster size around *n* = 32 parasites in four individual mice. Imaging of T cell clusters started at different times in individual mice and followed for different lengths of time. Note that as we observed before, 12 parasites had no T cells near them at both observations. Overall, there was a statistically significant but small change in the cluster size of the imaging period (as summarized in Figure 6B).

**Figure S3.**
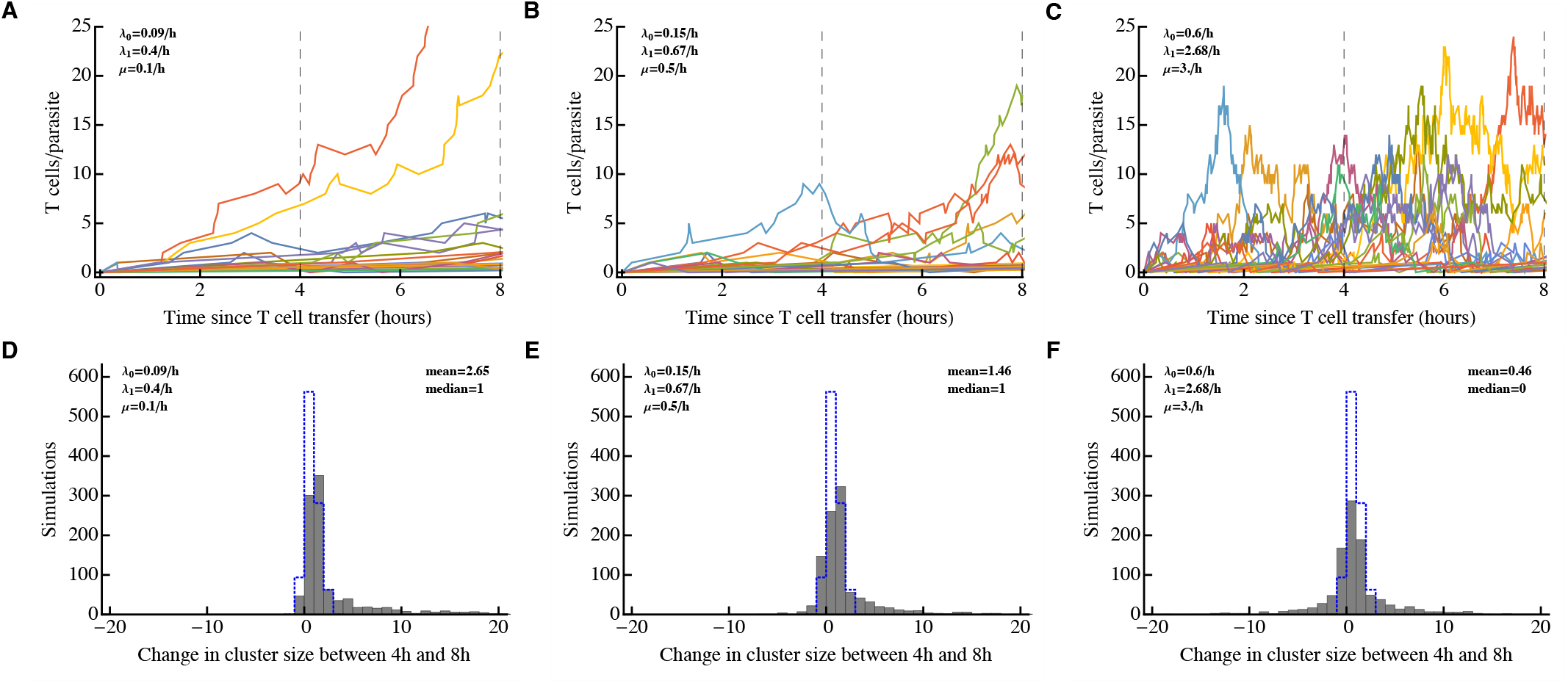
Stochastic simulations of cluster formation suggest an upper limit on the rate of T cell exit from the clusters. We ran Gillespie simulations of the cluster formation assuming different constant (time-independent) values for the entry rates into the cluster (λ_0_ and λ_1_) and exit rates from the cluster (*μ*) found by fitting the DD recruitment model to experimental data in Figure 6C-D. Three values of the exit rate were fixed: *μ* = 0.1/h (panels A&D), *μ* = 0.5/h (panels D&E), and *μ* = 3/h (panels C&F) and remaining parameters were estimated by fitting the model (eqns. (1)–(2)) to data (Figure 2B). These parameters are shown on individual panels. We simulated changes in cluster size for *n* = 10^3^ parasites. Panels A-C show sample trajectories of cluster sizes of 20 of such simulations, and panels D-F show the change in the size of the cluster between 4 and 8 hours after start of simulation for all simulations (solid bars) or changes in cluster sizes as was observed in experimental data (dotted bars, see also Figure 6B). These simulations indicate that at high exit rates (~ *μ* = 1 – 3/h) and at high entry rates there are large fluctuations in the cluster sizes between 4 and 8 hours (panels C&F) which is not observed in experimental data. Thus, in the 4-8 hour time period exit and entry rates cannot be extremely large for the DD recruitment model to be consistent with experimental data. Furthermore, simulations with smaller rates (panels A&D) also indicate increase in the average cluster size over time (since λ_1_ > *μ*) which is also not consistent with the change in cluster size at 4-8 hours post T cell transfer.

**Figure S4.**
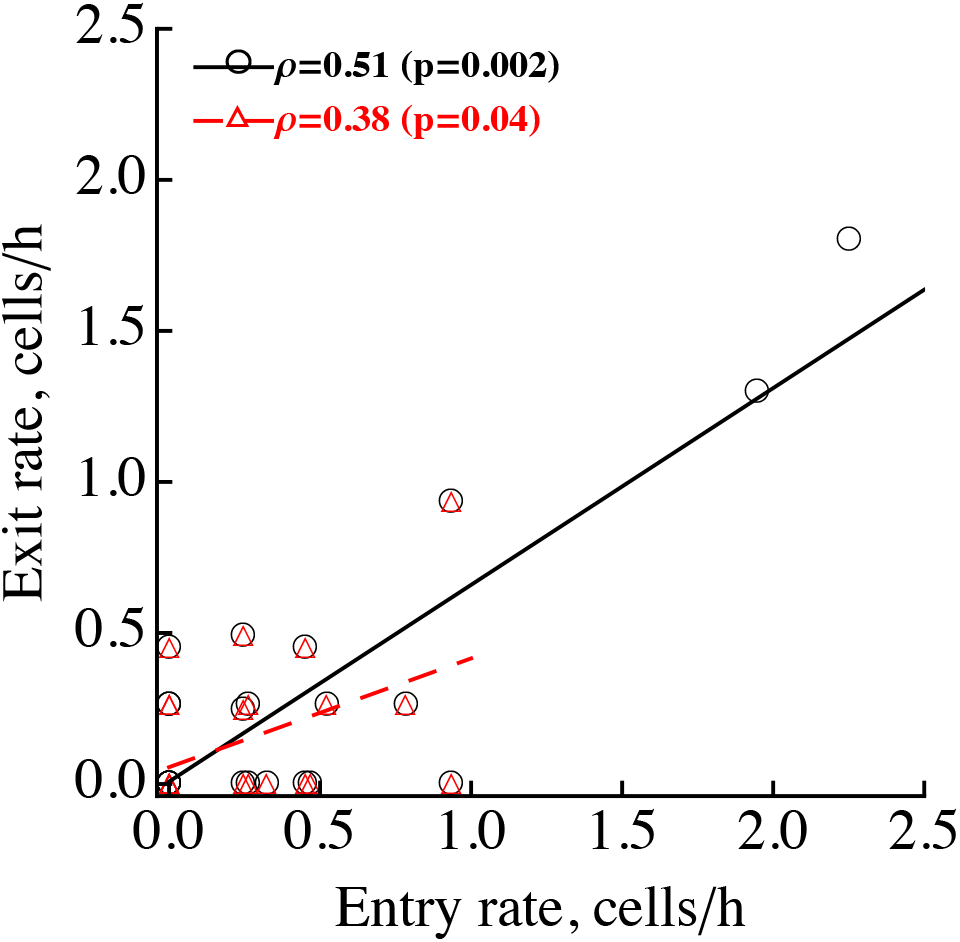
Experimentally measured rate of T cell exit from the cluster correlates with the rate of T cell entry into the cluster. We plotted the correlation between the experimentally measured number of T cells coming with a 40 *μ*m radius of a given parasite per unit of time (entry rate, see Figure 6E) and the number of T cells leaving a given cluster per unit of time (exit rate, see Figure 6F) for *n* = 32 parasites. P-values were calculated using Spearman Rank correlation test (with correlation coefficient *ρ* indicated), and lines indicate trends of the correlation found using a linear regression. The statistical significance of the correlation is shown for all data (circles) or for data that excluded two potential outliers (triangles).

